# Loss of Fas-signaling in pro-fibrotic fibroblasts impairs homeostatic fibrosis resolution and promotes persistent pulmonary fibrosis

**DOI:** 10.1101/2020.08.18.255869

**Authors:** Elizabeth F. Redente, Sangeeta Chakraborty, Satria Sajuthi, Bart P. Black, Benjamin L. Edelman, Max A. Seibold, David W.H. Riches

**Affiliations:** Program in Cell Biology, Department of Pediatrics and Denver CO 80206; Center for Genes, Environment and Health, National Jewish Health, Denver CO 80206; Department of Immunology and Microbiology, Aurora, CO 80010; Department of Pharmacology, Aurora, CO 80010; Division of Pulmonary Sciences and Critical Care Medicine, Department of Medicine, University of Colorado School of Medicine, Aurora, CO 80010; Department of Research, Veterans Affairs Eastern Colorado Health Care System, Denver, CO 80220

## Abstract

Idiopathic pulmonary fibrosis (IPF) is a progressive, irreversible fibrotic disease of the distal lung alveoli that culminates in respiratory failure and reduced lifespan. Unlike normal lung repair in response to injury, IPF is associated with the accumulation and persistence of fibroblasts and myofibroblasts and continued production of collagen and other extracellular matrix (ECM) components. Prior *in vitro* studies have led to the hypothesis that the development of resistance to Fas-induced apoptosis by lung fibroblasts and myofibroblasts contibributes to their accumulation in the distal lung tissues of IPF patients. Here, we test this hypothesis *in vivo* in the resolving model of bleomycin-induced pulmonary fibrosis in mice. Using genetic loss-of-function approaches to inhibit Fas signaling in fibroblasts, novel flow cytometry strategies to quantify lung fibroblast subsets and transcriptional profiling of lung fibroblasts by bulk and single cell RNA-sequencing, we show that Fas is necessary for lung fibroblast apoptosis during homeostatic resolution of bleomycin-induced pulmonary fibrosis *in vivo*. Furthermore, we show that loss of Fas signaling leads to the persistence and continued pro-fibrotic functions of lung fibroblasts. Our studies provide novel insights into the mechanisms that contribute to fibroblast survival, persistence and continued ECM deposition in the context of IPF and how failure to undergo Fas-induced apoptosis prevents fibrosis resolution.

## INTRODUCTION

Idiopathic pulmonary fibrosis (IPF), a progessive fibrotic lung disorder of unacceptably high morbidity and mortality, develops in response to initial alveolar epithelial injury and results in an abberant repair response in which resident lung interstitial and perivascular fibroblasts proliferate and persist within the collapsed alveoli and septal interstitium (1). These fibroblasts continue to produce excessive amounts of extrcellular matrix (ECM), which in turn leads to relentless fibrosis, progressive declines in gas exchange and eventual respiratory failure (1, 2). Acute respiratory distress syndrome (ARDS) patients also exhibit high morbidity and mortality resulting from initial alveolar epithelial injury (3). They also accumulate fibroblasts and myofibroblasts (collectively referred to herein as “pro-fibrotic fibroblasts”) that secrete collagen and ECM components within the injured alveoli, early in their clinical course. However, while both conditions share some similarities in the development of the early response to injury, the clinical outcomes of these two disorders are substantially different. IPF patients develop a persistent and progressive fibrosis considered by many to be irreversible. By contrast, while ARDS patients experience high initial mortality, the early fibroproliferative response can resolve and lung structure and function return towards normality in some survivors (4–7).

Little is known about the factors that distinguish these different outcomes in IPF patients and ARDS survivors. Fibrosis is generally considered to be detrimental for clinical outcomes. However, the early fibrosis that develops in ARDS patients could be viewed as a beneficial response in which the provisional ECM scaffold produced by proliferating pro-fibrotic fibroblasts supports the regenerating alveolar epithelium and endothelium. As alveolar structure and function are restored, the majority of the pro-fibrotic fibroblasts undergo apoptosis and clearance (8, 9). During this period of tissue regeneration, macrophages and other cells are thought to degrade and phagocytose the fibrotic provisional ECM scaffold through an integrated process that we define herein as “homeostatic fibrosis resolution”. In contrast to this trophic regenerative process, the persistent and progressive pulmonary fibrosis that defines IPF patients can be viewed as a failure in homeostatic fibrosis resolution. We hypothesize that a failure in pro-fibrotic lung fibroblast apoptosis and clearance plays a fundamentally important role in impeding homeostatic fibrosis resolution in IPF.

Previous studies by ourselves and others have provided insight into the mechanisms that control lung fibroblast apoptosis. In particular, susceptibility to apoptosis induction by the death receptor, Fas, has been shown to play a key role (10–15). Fas is expressed by lung fibroblasts and once a pre-defined threshold has been exceeded, Fas ligation induces apoptosis (11, 16). In contrast, lung fibroblasts from IPF patients exhibit variable resistance to Fas-induced apoptosis (12, 14) due to both down-regulation of Fas expression (16, 17) and increased expression of multiple anti-apoptotic genes, including Bcl-2, XIAP, cFLIP_L_ and PTPN13 (10, 18–20). Together, these *in vitro* studies have fueled the notion that the acquired resistance of lung fibroblasts to Fas-induced apoptosis promotes pro-fibrotic fibroblast accumulation in the persistently fibrotic lungs of IPF patients (10, 14, 15, 19, 21, 22). Until now, this assertion has remained unproven *in vivo*.

To address this knowledge gap, we genetically deleted Fas in fibroblasts and investigated the functional consequences on the normally resolving model of bleomycin-induced fibrosis. Intratracheal instillation of a single dose of bleomycin induces inflammatory epithelial injury followed by a resolving fibrotic response accompanied by regeneration of the distal lung architecture (8, 23). Thus, the bleomycin model recapitulates the acute lung injury, early fibrosis followed by fibrosis resolution and lung regeneration seen in ARDS survivors (3, 24), and represents an authentic model of homeostatic fibrosis resolution at later time points. As we will show, loss of Fas signaling in pro-fibrotic fibroblasts impairs homeostatic fibrosis resolution and results in fibroblast persistence in lung tissues along with persistent pulmonary fibrosis, replicating a critical feature of IPF. Furthermore, Col1a1- and αSMA-reporter mice and bulk and single cell RNA-sequencing assessments of purified lung fibroblasts showed that Fas deficiency in fibroblasts results in preserved patterns of increased ECM and fibrotic gene expression in the persistently fibrotic lungs. We conclude that Fas-induced fibroblast apoptosis plays a necessary role in fibroblast elimination during homeostatic fibroblast resolution, and that resistance to Fas-mediated apoptosis promotes persistent pulmonary fibrosis.

## RESULTS

### Fas deletion in fibroblasts impairs lung homeostatic fibrosis resolution

To investigate the functional role of Fas expression by fibroblasts in homeostatic fibrosis resolution, we deleted Fas in mesenchymal cells by breeding *Fas^fl/fl^* mice with *Dermo1(Twist2)-Cre* mice (25) to create *Dermo1-Cre;Fas^fl/fl^* mice (abbreviated as Dermo1-Cre;Fas^−/−^). To ensure that Fas was deleted, we isolated lung fibroblasts from wild type and Dermo1-Cre;Fas^−/−^ mice and assessed their level of cell surface Fas expression by flow cytometry, and their ability to undergo apoptosis: (i) following sensitization with TNF-α and IFN-α and Fas-ligation with agonistic anti-Fas antibody (Jo2), and (ii) in response to staurosporine, an activator of the intrinsic apoptosis pathway. In comparison to wild type lung fibroblasts, fibroblasts from Dermo1-Cre;Fas^−/−^ mice displayed minimal cell surface Fas basally or in response to stimulation with TNF-α and IFN-α (**Fig. 1A**) and were not susceptible to Fas-induced apoptosis (**Fig. 1B**). However, fibroblasts from both wild type and Dermo1-Cre;Fas^−/−^ mice showed similar levels of apoptosis induction in response to staurosporine (**Fig. 1B**). Together, these data indicate that *in vivo* genetic deletion of Fas in mesenchymal cells led to loss of cell surface Fas and corresponding apoptosis induction by the Fas-induced extrinsic apoptosis pathway, but not by staurosporine.

**Figure 1.**
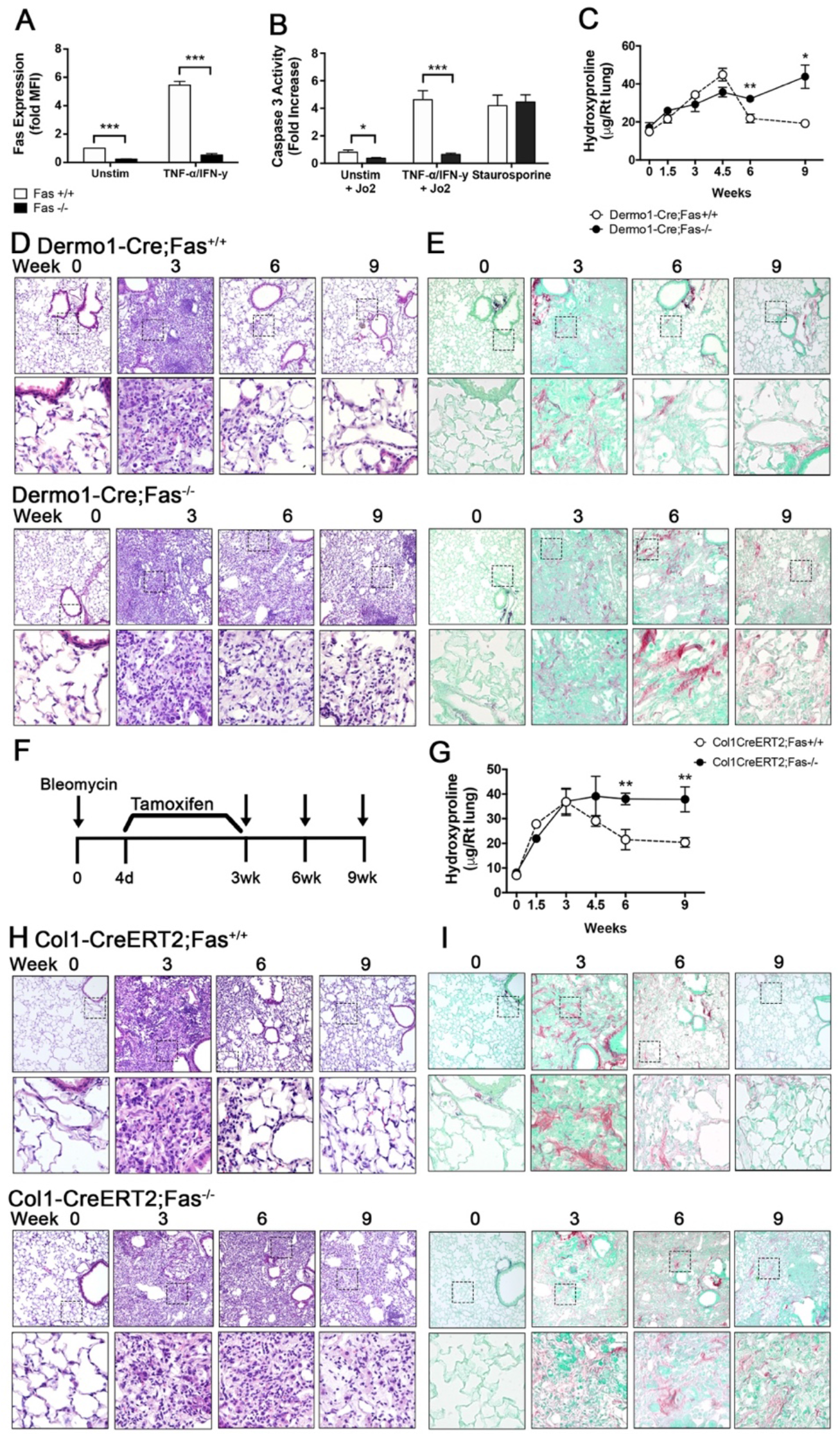
Fas expression in mesenchymal cells is essential for their spontaneous apoptosis during fibrosis resolution. (A) Cell surface expression on cultured fibroblasts from wild type and Fas-deficient mice. (B) Caspase 3 activity in wild type and Fas-deficient fibroblasts after Fas-induced apoptosis. (C) Hydroxyproline levels in the lungs over time after bleomycin in wild type and Dermo1-Cre;Fas^−/−^ mice. (D-E) Representative H&E and Picrosirius Red staining lung sections over time in wild type and Dermo1-Cre;Fas^−/−^ mice. (F) Sketch illustrating instillation time, tamoxifen dosing and harvest time points in Col1-CreERT2;Fas^fl/fl^ mice. (G) Hydroxyproline levels in the lungs over time after bleomycin in Col1-CreERT2;Fas^+/+^ and Col1-CreERT2;Fas^−/−^ mice. (H-I) Representative H&E and Picrosirius Red staining lung sections over time in Col1-CreERT2;Fas^+/+^ and Col1-CreERT2;Fas^−/−^ mice. (mean +/− SEM, n=6-8). *p<0.05, **p<0.01, Student’s t test. Upper histology panels 20X, lower panels 40X.

Next, we evaluated the ability of Dermo1-Cre;Fas^−/−^ mice to undergo homeostatic fibrosis resolution in response to a single intratracheal instillation of bleomycin (1.5 U/kg). Wild type mice exhibit a predictable course of acute inflammatory lung injury at 1 week, fibrosis development at 2-4 weeks and resolution to near normal lung structure and function by 6-9 weeks (23). Dermo1-Cre;Fas^−/−^ and control Dermo1-Cre;Fas^+/+^ mice developed similar amounts of bleomycin-induced pulmonary fibrosis, as reflected by lung hydroxyproline levels at 1.5, 3 and 4.5 weeks (**Fig. 1C**). Both strains also exhibited similar changes in lung histopathology and patterns of peribronchiolar and parenchymal collagen deposition as revealed by Picrosirius red staining (**Figs. 1D,E**). However, unlike Dermo1-Cre;Fas^+/+^ mice, which had undergone homeostatic fibrosis resolution at 6 and 9 weeks, the lungs of Dermo1-Cre;Fas^−/−^ mice showed persistent fibrosis and histopathologic distortion at 4.5, 6 and 9 (**Figs. 1C,D,E**). Wild type C57Bl/6 mice and Dermo1-Cre;Fas^+/+^ mice showed identical patterns of fibrosis development and homeostatic fibrosis resolution (not shown).

Pro-fibrotic fibroblasts express multiple ECM genes, including Col1a1 (**Fig. S1**). We therefore determined if conditional deletion of Fas in Col1a1-expressing cells phenocopied the impairment of homeostatic fibrosis resolution seen in Dermo1-Cre;Fas^−/−^ mice. *Fas^fl/fl^* mice were bred with *Col1a1-CreERT2* mice (26) to create *Col1a1-Cre-ERT2;Fas^fl/fl^* mice, which we refer to as Col1-CreERT2;Fas^−/−^ following tamoxifen injection, and Col1-CreERT2;Fas^+/+^ following corn oil vehicle injection. As illustrated in **Fig. 1F**, *Col1-CreERT2;Fas^fl/fl^* mice were instilled with bleomycin, treated with either tamoxifen or corn oil twice weekly during fibrosis development between weeks 0.5-3, and assessed for fibrotic outcomes between weeks 1.5 and 9. Recombination at the Fas^fl/fl^ locus in tamoxifen-treated mice was confirmed by PCR analysis of tail tip fibroblast DNA (**Fig. S2**). **Fig. 1G** shows that tamoxifen-injected Col1-CreERT2;Fas^−/−^ mice and corn oil injected Col1-CreERT2;Fas^+/+^ mice developed similar amounts of fibrosis at 1.5 and 3 weeks. However, whereas bleomycin-instilled, Col1-CreERT2;Fas^+/+^ mice underwent homeostatic fibrosis resolution between 4.5 and 9 weeks, homeostatic fibrosis resolution was impaired in the lungs of bleomycin-instilled Col1-CreERT2;Fas^−/−^ mice, as reflected by persistently elevated lung hydroxyproline levels (**Fig. 1G**) and sustained fibrotic lung histology patterns (**Figs. 1H,I**). Wild type C57Bl/6 mice and Col1-CreERT2;Fas^+/+^ mice showed indistinguishable patterns of fibrosis development and homeostatic fibrosis resolution ((23) and not shown). Together, these data show that loss of Fas-signaling in Col1a1-expressing pro-fibrotic fibroblasts also led to impaired homeostatic fibrosis resolution and persistent pulmonary fibrosis.

### Fas deletion in fibroblasts promotes lung fibroblast persistence

We next assessed the consequences of loss of Fas-signaling on lung fibroblast numbers. Dermo1-Cre;Fas^−/−^ and Dermo1-Cre;Fas^+/+^mice were instilled with saline or bleomycin and lung tissues harvested at 3, 6 and 9 weeks. Lung fibroblasts were initially identified in lung paraffin sections by immunofluorescent staining for αSMA and S100A4 (FSP1). **Fig. 2A** shows that while naïve lung tissues exhibited minimal staining for S100A4 and αSMA in the distal alveoli, bleomycin-instilled Dermo1-Cre;Fas^−/−^ mice and Dermo1-Cre;Fas^+/+^ mice exhibited abundant and concordant staining of αSMA+, S100A4+ and αSMA+S100A4+ fibroblasts in the fibrotic lung parenchyma at 3 weeks. However, whereas parenchymal αSMA and S100A4 staining was reduced at 6 and 9 weeks in the regenerated lung parenchyma of Dermo1-Cre;Fas^+/+^ mice, αSMA+, S100A4+ and αSMA+S100A4+ fibroblasts remained in the persistently fibrotic lungs of Dermo1-Cre;Fas^−/−^ mice at 6 and 9 weeks (**Fig. 2A**).

**Figure 2.**
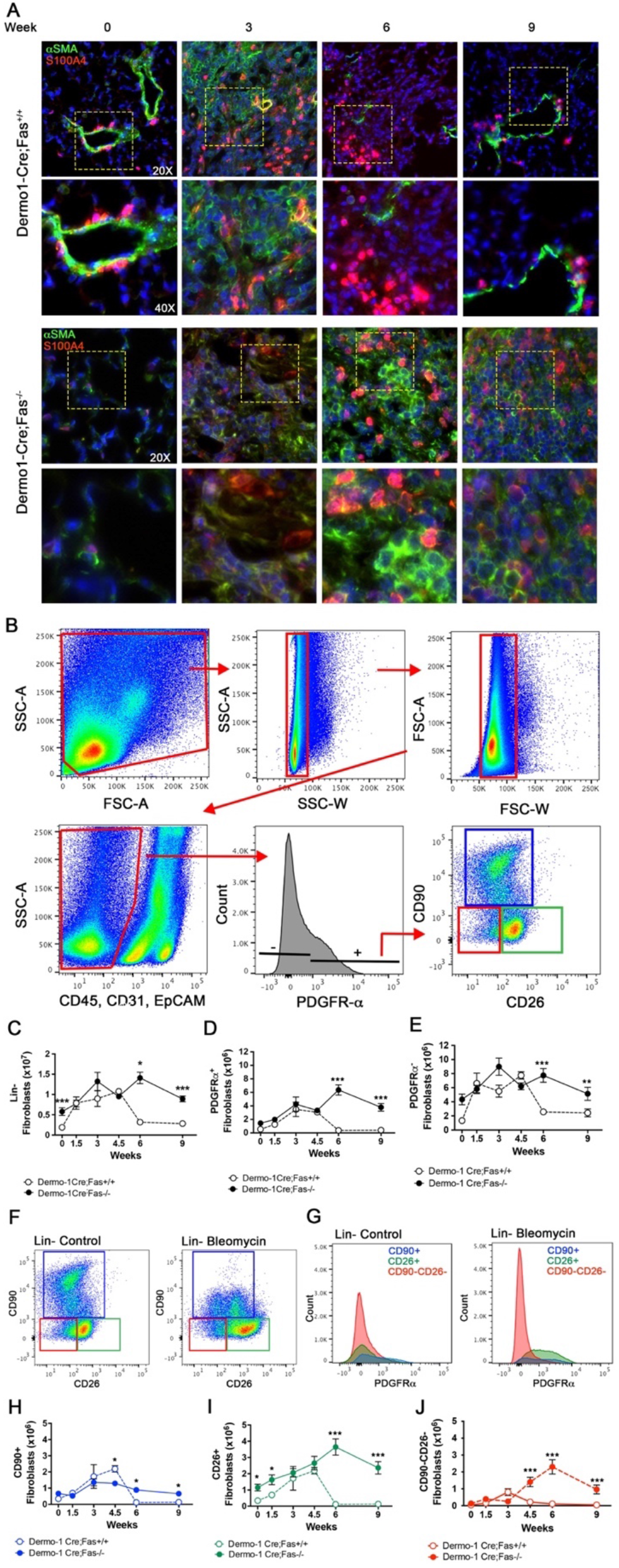
Fibroblasts persist in fibrotic lungs in the absence of Fas. (A) αSMA (green) and S100A4 (red) immunofluorescent staining of lungs over time after bleomycin in Dermo1-Cre;Fas^+/+^ and Dermo1-Cre;Fas−/− mice. (B) Flow cytometry strategy to identify fibroblast subsets. (C-E) Quantitation of Lin−, PDGFRα+ and PDGFRα- fibroblasts populations. (F-G) Representative gating strategy to identify Lin-subsets: CD90+CD26-, CD26+CD90- and CD90-CD26- and their relative expression of PDGFRα in control and after bleomycin treatment. (H-J) Quantitation of CD90+CD26-, CD26+CD90- and CD90-CD26-fibroblast subsets. (mean +/− SEM, n=6-8). *p<0.05, **p<0.01, ***p<0.001 Student’s t test. Upper panels 20X, lower panels 40X.

To confirm these findings, we used enzymatic lung cell dispersal and multi-parameter flow cytometry to assess the consequences of loss of Fas-signaling on lung fibroblast numbers in Dermo1-Cre-expressing cells. Lung single cell suspensions were analyzed using the gating strategy shown in **Fig. 2B.** From these data we quantified the number of lineage negative (Lin-) cells, which are throught to be primarily fibroblasts (27–29), by excluding EpCAM/CD326+ (epithelial), CD45+ (hematopoietic) and CD31+ (endothelial) positive populations. Dermo1-Cre;Fas^+/+^ and Dermo1-Cre;Fas^−/−^ mice were instilled with saline or bleomycin and the numbers of Lin-cells quantified for up to 9 weeks. **Fig. 2C** shows that during the fibrosis development between 1.5-3 weeks, the number of Lin-lung fibroblasts increased ~2.3-fold, with similar numbers being seen in both Dermo1-Cre;Fas^+/+^ and Dermo1-Cre;Fas^−/−^ mice. Within the Lin-fraction, approximately 33% expressed variable levels of the pan-fibroblast marker PDGFRα while 66% were PDGFRα. **Fig. 2C,D,E** also shows that whereas the numbers of total lung Lin-cells, Lin-PDGFRα+ and Lin-PDGFRα-cells in Dermo1-Cre;Fas^+/+^ were progressively reduced by 6 and 9 weeks, their numbers remained elevated at 6 and 9 weeks in Dermo1-Cre;Fas^−/−^.

Novel stratiegies to idenfity pulmonary fibroblast subsets by flow cytometry are still needed. Therefore, we sought to utilize cell surface markers to idenfity fibroblasts and segregate them into function-related subsets using PDGFRα (a pan fibroblast marker), CD90 (Thy1) and CD26 (Dpp4). CD90 (Thy1) is expressed by lipid-laden resident alveolar interstitial fibroblasts and perivascular pericyte-like cells (30–33); while CD26 is expressed by matrix-producing reticular dermal fibroblasts (34). We confirmed the presence of CD90 and CD26 in lung fibroblasts by flow cytometry of dispersed lung tissues from saline and bleomycin-instilled C57Bl/6 mice at 3 weeks (**Fig. S3**) (27, 35–37). Three Lin-subsets which express variable levels of PDGFRα were identified: CD90+CD26-, CD90-CD26+ and CD90-CD26-, which we abbreviate as CD90+, CD26+ and CD90-CD26-subsets (**Fig. 2F,G**). CD90+ fibroblasts were the most abundant subset in the lungs of naïve and saline-instilled C57Bl/6 mice in comparison to CD26+ cells and CD90-CD26-cells. (**Fig. S3**). However, whereas the numbers of CD90+ fibroblasts were only modestly increased (1.2-fold) following bleomycin instillation, CD26+ and CD90-CD26-fibroblasts underwent a large expansion (7.7-fold and 9.6-fold, respectively) compared to saline-instilled mice at 3 weeks (**Fig. S3**).

We next assessed the consequences of Fas deletion in fibroblasts on the numbers of CD90+, CD26+ and CD90-CD26-fibroblasts in bleomycin-instilled Dermo1-Cre;Fas^−/−^ and Dermo1-Cre;Fas^+/+^ mice. **Fig. 2H,I,J** shows that following bleomycin instillation, the numbers of CD90+, CD26+ and CD90-CD26-fibroblasts expanded as fibrosis developed in Dermo1-Cre;Fas^+/+^ mice, peaking between 1.5 and 4.5 weeks before declining back to baseline as the mice underwent homeostatic fibrosis resolution by 6 and 9 weeks. Interestingly, the decline in fibroblast numbers was slightly delayed when compared to the rate of decline in lung collagen levels (**Fig. 1B,C,D**). Lung CD90+, CD26+ and CD90-CD26-fibroblast numbers also increased by 3 weeks in Dermo1-Cre;Fas^−/−^ mice (**Fig. 2H,I,J**). However, in contrast to Dermo1-Cre;Fas^+/+^ mice, their numbers remained elevated at 4.5, 6 and 9 weeks in the lungs of bleomycin-instilled Col1-CreERT2;Fas^−/−^ mice (**Fig. 2H,I,J**).

To confirm these findings, *Col1-CreERT2;Fas^fl/fl^* mice were instilled with saline or bleomycin, injected with tamoxifen or corn oil between weeks 0.5-3 (**Fig. 1F**) and lung tissues harvested at 3, 6 and 9 weeks. As was seen with the Dermo1-Cre;Fas^+/+^ and Dermo1-Cre;Fas^−/−^, bleomycin-instilled Col1-CreERT2;Fas^−/−^ mice and Col1-CreERT2;Fas^+/+^ mice exhibited similar staining of αSMA+, S100A4+ and αSMA+S100A4+ fibroblasts at 3 weeks. However, whereas peripheral αSMA and S100A4 staining was reduced at 6 and 9 weeks in the regenerated lung parenchyma of Col1-CreERT2;Fas^+/+^ mice, αSMA+, S100A4+ and αSMA+S100A4+ fibroblasts were still detected in lungs of Col1-CreERT2;Fas^−/−^ mice at 6 and 9 weeks (**Fig. 3A**). Similarly, flow cytometry revealed a similar ~2-fold increase in Lin-fibroblast numbers in both Col1-CreERT2;Fas^−/−^ and Col1-CreERT2;Fas^+/+^ mice during fibrosis development between 1.5-3 weeks and within the Lin-fraction, approximately 35% expressed variable levels of PDGFRα while 65% were PDGFRα-. **Fig. 3B,C,D** also shows that whereas the numbers of total lung Lin-cells, Lin-PDGFRα+ and Lin-PDGFRα-cells in Col1-CreERT2;Fas^+/+^ declined towards baseline by 6 and 9 weeks, their numbers remained elevated at 6 and 9 weeks in Col1-CreERT2;Fas^−/−^ mice. Lung CD90+, CD26+ and CD90-CD26-fibroblast numbers also increased between 1.5-4.5 weeks in Col1-CreERT2;Fas^−/−^ mice (**Fig. 3E,F,G,H,I**) but, in contrast to Col1-CreERT2;Fas^+/+^ mice, their numbers remained significantly elevated at 6 and 9 weeks (**Fig. 2G,H,I**). To affirm the specificity of the Col1a1-promoter driven Fas-deletion strategy, we lineage traced with TdTomato. **Fig. S4** shows that TdTomato was primarily expressed in Lin-PDGFRα+ cells. Thus, Fas deficiency in Dermo1-Cre and Col1a1-expressing fibroblasts impeded fibroblast elimination from lung tissues during homeostatic resolution of bleomycin-induced fibrosis.

**Figure 3.**
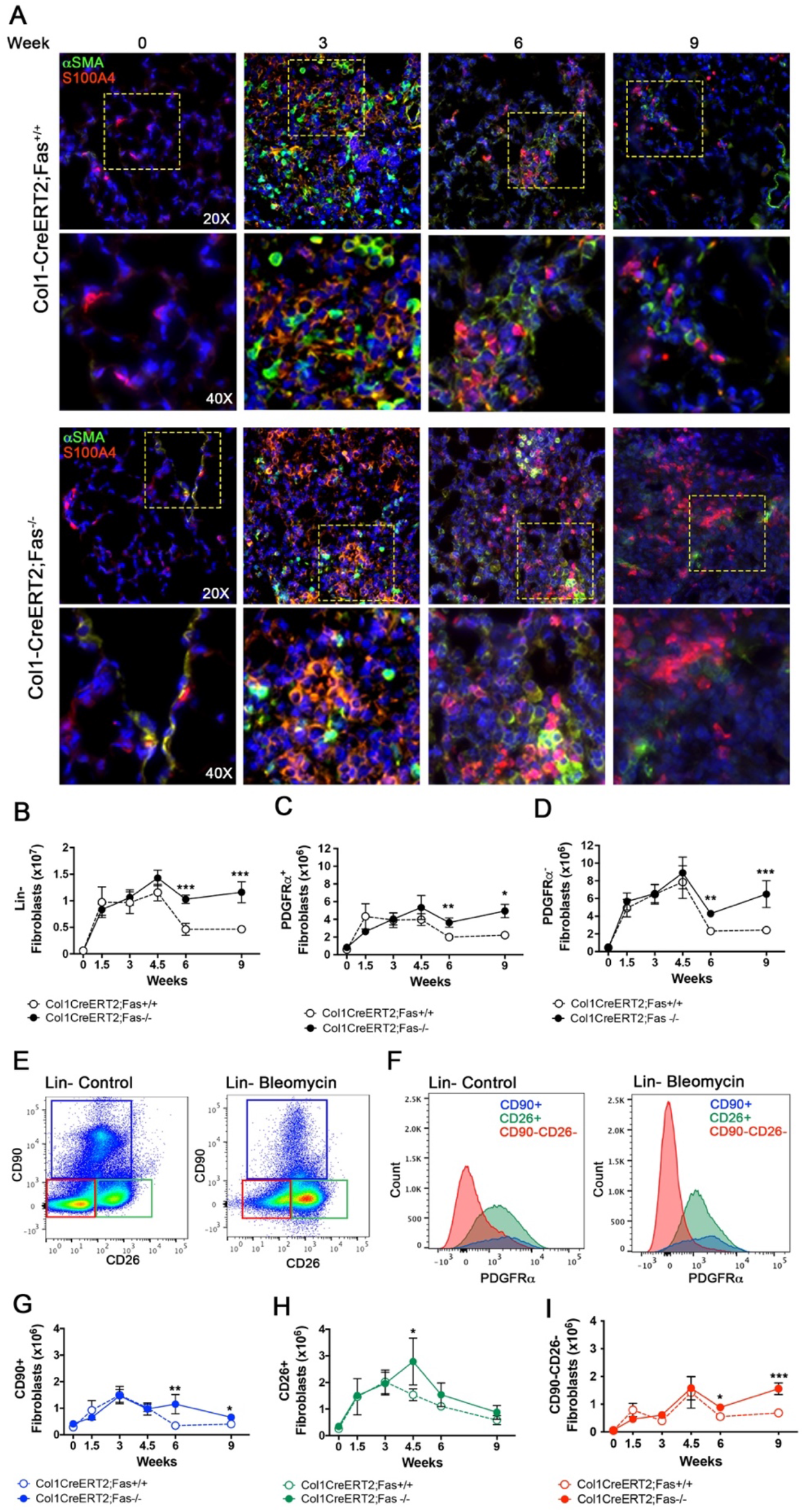
Deletion of Fas in Col-1 fibroblasts permits their persistence during fibrosis. αSMA (green) and S100A4 (red) immunofluorescent staining of lungs over time after bleomycin in Col1-CreERT2;Fas^+/+^ and Col1-CreERT2;Fas^−/−^ mice. (B-D) Quantitation of Lin-, PDGFR α+ and PDGFR α-fibroblasts populations. (E-F) Representative gating strategy to identify Lin-subsets: CD90+CD26-, CD26+CD90- and CD90-CD26- and their relative expression of PDGFRα in control and after bleomycin treatment. (G-I) Quantitation of CD90+CD26-, CD26+CD90- and CD90-CD26-fibroblast subsets. (mean +/− SEM, n=6-9). *p<0.05, **p<0.01, Student’s t test. Upper panels 20X, lower panels 40X.

### Fas deletion in fibroblasts impairs their apoptosis during homeostatic fibrosis resolution

We next investigated the effect of Fas-deficiency on lung fibroblast apoptosis *in vivo*. Dermo1-Cre;Fas^+/+^ and Dermo1-Cre;Fas^−/−^ mice were instilled with saline or bleomycin and lung tissues harvested from 1.5-9 weeks. Lung sections were then subjected to triple-label immunofluorescent staining for TUNEL, α-SMA and S100A4 (**Fig. 4A**) and TUNEL+ fibroblasts were quantified. **Fig. 4B** shows minimal numbers of apoptotic TUNEL+ fibroblasts in naïve Dermo1-Cre;Fas^+/+^. Increased numbers were first detected in bleomycin-instilled mice at 1.5 weeks, peaked at 3 weeks and declined back to naïve numbers at 6 weeks, confirming a previous report (9). Following Fas deletion in fibroblasts in Dermo1-Cre;Fas^−/−^ mice, the number of TUNEL+ fibroblasts were not significantly elevated over baseline values at any time point, but when compared to Dermo1-Cre;Fas^+/+^ mice, were significantly reduced at 1.5 weeks (p=0.06), 3 weeks (p<0.05) and 4.5 weeks (p=0.08) (**Fig. 4A, B**). We confirmed these data in naïve and bleomycin-instilled *Col1-CreERT2;Fas^fl/fl^* mice following tamoxifen or corn oil injection between 0.5-3 weeks, with harvest and analysis for TUNEL+ cells for up to 9 weeks. **Fig. 4C,D** shows that Col1-CreERT2;Fas^+/+^ mice also exhibited a peak of TUNEL+ apoptotic lung fibroblasts at 3 weeks, while TUNEL+ fibroblast numbers were not elevated above baseline at any time point in Col1-CreERT2;Fas^−/−^ mice (0=0.02). Taken together, these data show that Fas deletion in fibroblasts prevents homeostatic fibroblast apoptosis in bleomycin-injured fibrotic lung tissues.

**Figure 4.**
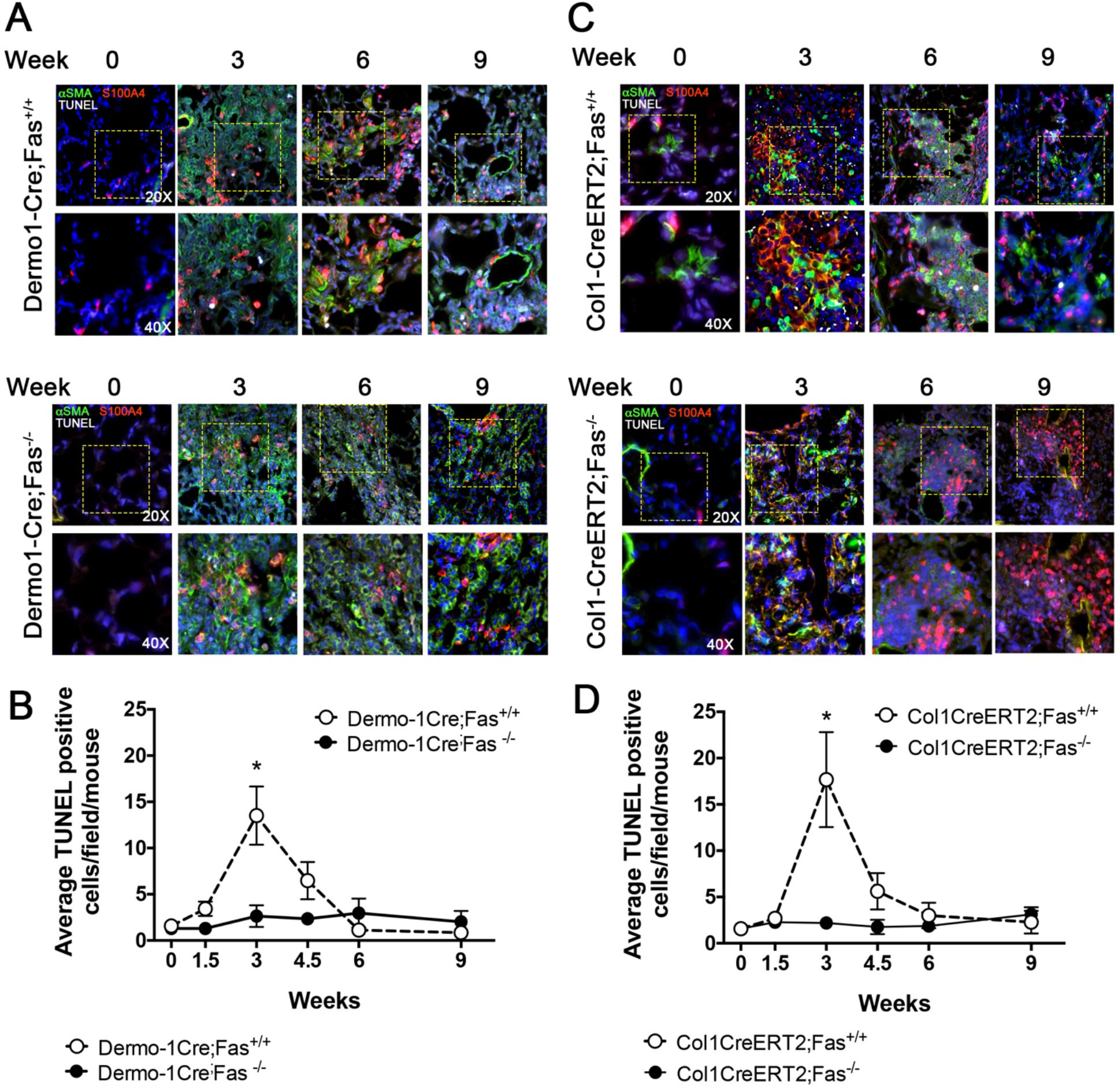
Loss of Fas reduces apoptosis of fibroblasts during fibrosis resolution in vivo. (A-B) Immunofluorescent staining and quantitation of fibroblasts using antibodies for αSMA (green), S100A4 (red) and TUNEL (white) in Dermo1-Cre;Fas^+/+^ and Dermo1-Cre;Fas−/− mice over time after bleomycin. (C-D) Immunofluorescent staining and quantitation of fibroblasts using antibodies for αSMA (green), S100A4 (red) and TUNEL (white) in Col1-CreERT2;Fas^+/+^ and Col1-CreERT2;Fas−/− mice over time after bleomycin. (mean +/− SEM, n=10 images per animal/time point). *p<0.05, Student’s t test. Upper panels 20X, lower panels 40X.

### Fas-deficiency in Col1a1-expressing cells leads to persistent Col1a1 and α-SMA promoter activity

Col1-GFP mice express GFP under the control of the Col1a1 promoter (38); while SMA-RFP mice express RFP under the control of the α-SMA promoter (39). We used these mice to determine if the Fas-deficient fibroblasts that remain in persistently fibrotic lung tissues of bleomycin-instilled mice continue to functionally express these pro-fibrotic genes. Frozen lung sections from naïve mice revealed modest numbers of dim GFP+ fibroblastic interstitial cells (**Fig. 5A)**, while bleomycin-instilled Col1-GFP mice exhibited increased numbers and intensity of GFP+ interstitial cells peaking at 3 weeks before returning towards baseline by 6-9 weeks. Flow cytometry analysis of lung single cell suspensions from bleomycin-instilled Col1-GFP mice showed that Lin-PDGFRα+ CD90+, CD26+ and CD90-CD26-fibroblast subsets all exhibited increased GFP expression that peaked at 3 weeks before declining to baseline by 6 weeks (**Fig. 5B,C,D and Fig. S5**). Naïve SMA-RFP mice also exhibited minimal RFP expression in the distal lung, though the smooth muscle surrounding arteries and larger airways exhibited well-defined RFP fluorescence (**Fig. 5E)**. Bleomycin instillation resulted in increased numbers of interstitial cells expressing RFP in the fibrotic distal lung tissues in lung frozen sections (**Fig. 5E**) and increased numbers of RFP+Lin-PDGFRα+ CD90+, CD26+ and CD90-CD26-fibroblast subsets that peaked at 3 weeks before declining to baseline by 6 weeks (**Fig. 5F,G,H and Fig. S6**). Thus, Col1-GFP and SMA-RFP mice quantitatively report changes in pro-fibrotic fibroblast Col1a1 and αSMA-promoter activity, as previously reported (39–43).

**Figure 5.**
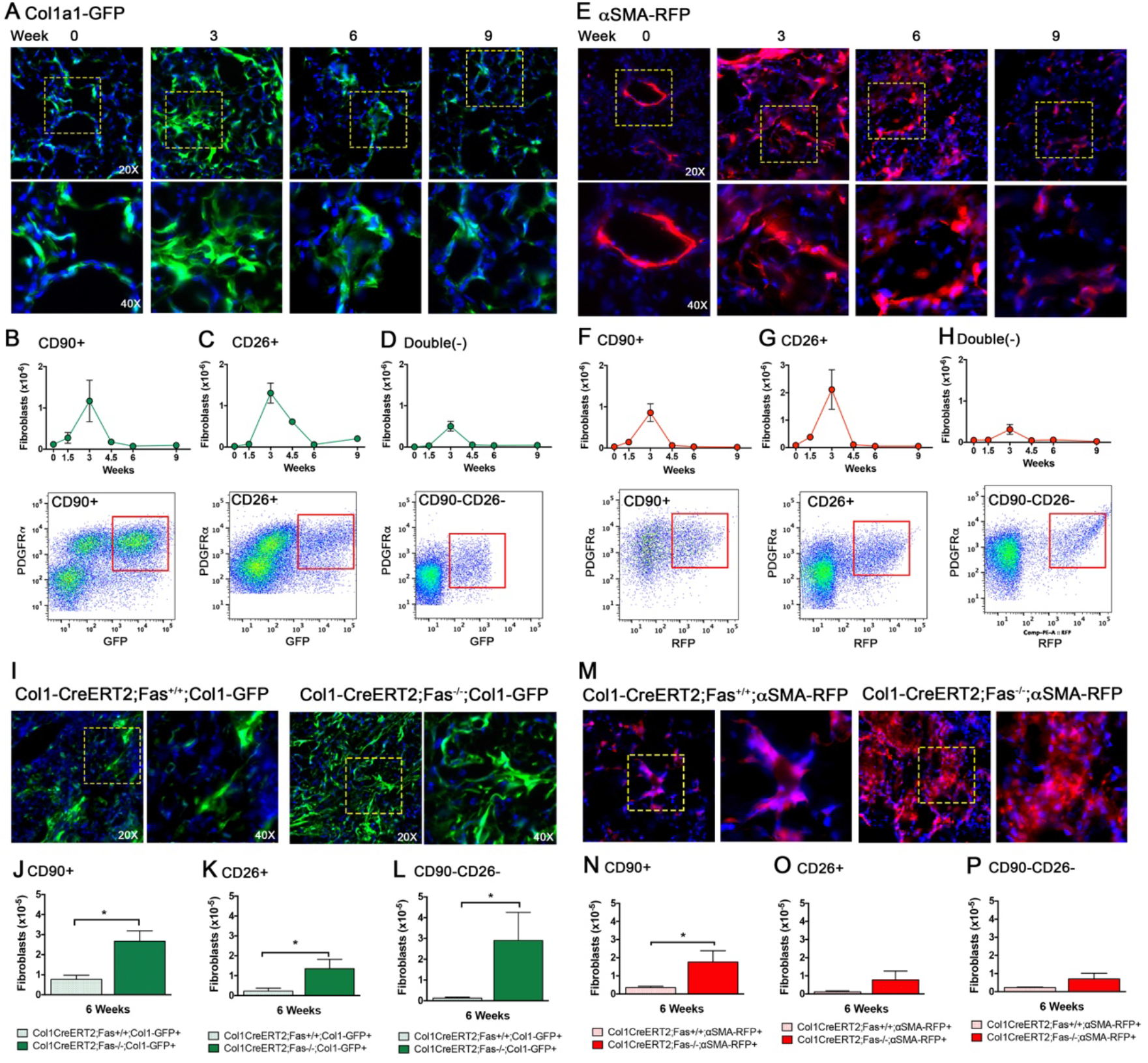
Fibroblast expression of Col1 and αSMA during fibrosis development, resolution and persistence. (A) GFP expression of Col1a1 in fibroblasts over time after bleomycin in Col1-GFP mice. (B-D) Quantitation and representative flow cytometry plots of GFP positive CD90+CD26-, CD26+CD90- and CD90-CD26-fibroblast subsets over time. (E) RFP expression of αSMA in fibroblasts over time after bleomycin in αSMA-RFP mice. (F-H) Quantitation and representative flow cytometry plots of RPF positive CD90+CD26-, CD26+CD90- and CD90-CD26-fibroblast subsets over time. (I) GFP expression of Col1a1 in fibroblasts 6 weeks after bleomycin in Col1-CreERT2;Fas^+/+^;Col1-GFP and Col1-CreERT2;Fas^−/−^;Col1-GFP mice. (J-L) Quantitation of GPF positive CD90+CD26-, CD26+CD90- and CD90-CD26-fibroblast subsets. (M) RFP expression of αSNA in fibroblasts 6 weeks after bleomycin in Col1-CreERT2;Fas^+/+^;αSMA-RFP and Col1-CreERT2;Fas^−/−^;αSMA-RFP mice. (N-P) Quantitation of RPF positive CD90+CD26-, CD26+CD90- and CD90-CD26-fibroblast subsets. (mean +/− SEM, n=5-12). *p<0.05, Student’s t test. Upper panels 20X, lower panels 40X.

To determine if the Fas-deficient fibroblasts that persist in fibrotic lung tissues continue to exhibit Col1a1 promoter activity, we bred *Col1-CreERT2;Fas^fl/fl^* mice with Col1-GFP mice to obtain *Col1-CreER;Fas^fl-fl^;Col1-GFP* mice. *Col1-CreER;Fas^fl-fl^;Col1-GFP* were instilled with bleomycin and injected with tamoxifen or corn oil between 0.5-3 weeks to yield Col1-CreERT2;Fas^−/−^;Col1-GFP and Col1-CreERT2;Fas^+/+^;Col1-GFP mice, respectively. The mice were harvested at 6 weeks and analyzed for the presence of GFP in lung fibroblasts in lung frozen sections and by multi-parameter flow cytometry. **Fig. 5I** shows that whereas GFP+ fibroblast numbers in frozen sections of bleomycin-instilled Col1-CreERT2;Fas^+/+^;Col1-GFP mice had returned to naïve levels at 6 weeks, increased numbers of GFP+ cells remained in the lungs of Col1-CreERT2;Fas^−/−^;Col1-GFP. Similarly, flow cytometry of lung single cell suspension revealed increased numbers of GFP+ Lin-PRGFRα+ CD90+, CD26+ and CD90-CD26-fibroblasts in the fibrotic lung tissues of Col1-CreERT2;Fas^−/−^;Col1-GFP mice at 6 weeks compared to bleomycin-instilled Col1-CreERT2;Fas^+/+^;Col1-GFP mice (**Fig. 5J,K,L**).

We also determined if the Fas-deficient fibroblasts that persist in fibrotic lungs also continued to express α-SMA by breeding *Col1-CreERT2;Fas^fl/fl^* mice with SMA-RFP reporter mice using the same strategy as discussed above to generate Col1-CreERT2;Fas^+/+^;SMA-RFP mice and Col1-CreERT2;Fas^−/−^;SMA-RFP mice. Compared to Fas-sufficient Col1-CreERT2;Fas^+/+^;SMA-RFP mice, lung tissues of Col1-CreERT2;Fas^−/−^; SMA-RFP mice displayed abundant RFP fluorescence in the fibroblastic cells in the persistently fibrotic distal lung tissues at 6 weeks (**Fig. 5M)**. Interestingly, and in contrast to the Col1-CreERT2;Fas^−/−^;Col1-GFP mice, flow cytometry of lung cell suspensions revealed significantly increased numbers of RFP+Lin-PRGFRα+ CD90+, but not of CD26+ and CD90-CD26-fibroblasts in the fibrotic lung tissues of Col1-CreERT2;Fas^−/−^;SMA-RFP mice (**Fig. 5N,O,P**). Neither GFP or RFP were detected in CD45+, CD31+ or CD326+ cells in bleomycin-instilled Col1-CreERT2;Fas^−/−^;Col1-GFP or Col1-CreERT2;Fas^−/−^;SMA-RFP mice, respectively (**Fig. S7**). Taken together, these data suggest that the Fas-deficient fibroblasts that remain in fibrotic lung tissues of bleomycin-instilled mice at 6 weeks continue to functionally express pro-fibrotic Col1a1- and αSMA-promoter activity.

### Bulk and single cell RNA-sequencing defines preserved pro-fibrotic fibroblast clusters and signatures in the absence of Fas-signaling

To further characterize the fibroblasts that persist in fibrotic lung tissues in the absence of Fas signaling, we conducted bulk and single cell RNA-sequencing (scRNA-seq) on sorted Lin-cells isolated from bleomycin-instilled Col1-CreERT2;Fas^−/−^ and Col1-CreERT2;Fas^+/+^ mice at 3 and 6 weeks. We also sorted Lin-cells from naïve Col1-CreERT2;Fas^fl/fl^ mice for comparison. Bulk RNA-seq showed that compared to naïve mice, bleomycin-instilled mice of both genotypes showed an expected increase in pro-fibrotic gene expression and fibrosis-associated pathways at 3 weeks (not shown). Principal Component Analysis (PCA) of highly variable genes partitioned samples by time of harvest after bleomycin instillation along the PC1 axis (30% variance) and by genotypic Fas deficiency along the PC4 axis (5% variance) (**Fig. 6A**). There were no significant differences in pro-fibrotic gene signatures in Lin-cells isolated from bleomycin-instilled Col1-CreERT2;Fas^−/−^ and Col1-CreERT2;Fas^+/+^ mice at 3 weeks. Pathway analysis of 393 upregulated differentially expressed genes (DEGs) at 6 weeks revealed that the most significantly enriched pathways in Col1-CreERT2;Fas^−/−^ mice were associated with extracellular matrix organization (p=6.32E-13), collagen organization (p=6.59E-09), endodermal cell differentiation (p=1.45E-05) and extracellular matrix-receptor interaction (p=0.0014) when compared to Col1-CreERT2;Fas^+/+^ mice (**Fig. 6B, Fig. S8 and Table S1**). The transcripts that contributed to these pathways included the typical pro-fibrotic genes Col1a1, Col5a1, Col6a1, Col7a1, Col8a1, Col11a1, Col12a1, Eln and Loxl2 (**Fig. 6C**). By contrast, Lin-cells from the Col1-CreERT2;Fas^+/+^ mice, which were undergoing homeostatic fibrosis resolution at 6 weeks, showed enrichment for regulation of cell migration (p=0.0079) and included genes involved in wound healing (Epb41l4b, Erbb3, Itgb3, Wnt7A and Nrg1), and Wnt signaling (Wnt10B, Fzd5, Wnt3A, Wnt7A, Sost, Rspo4, Wnt4) (**Fig. 6B**) (**Fig. 6B, Fig. S8 and Table S1**).

**Figure 6.**
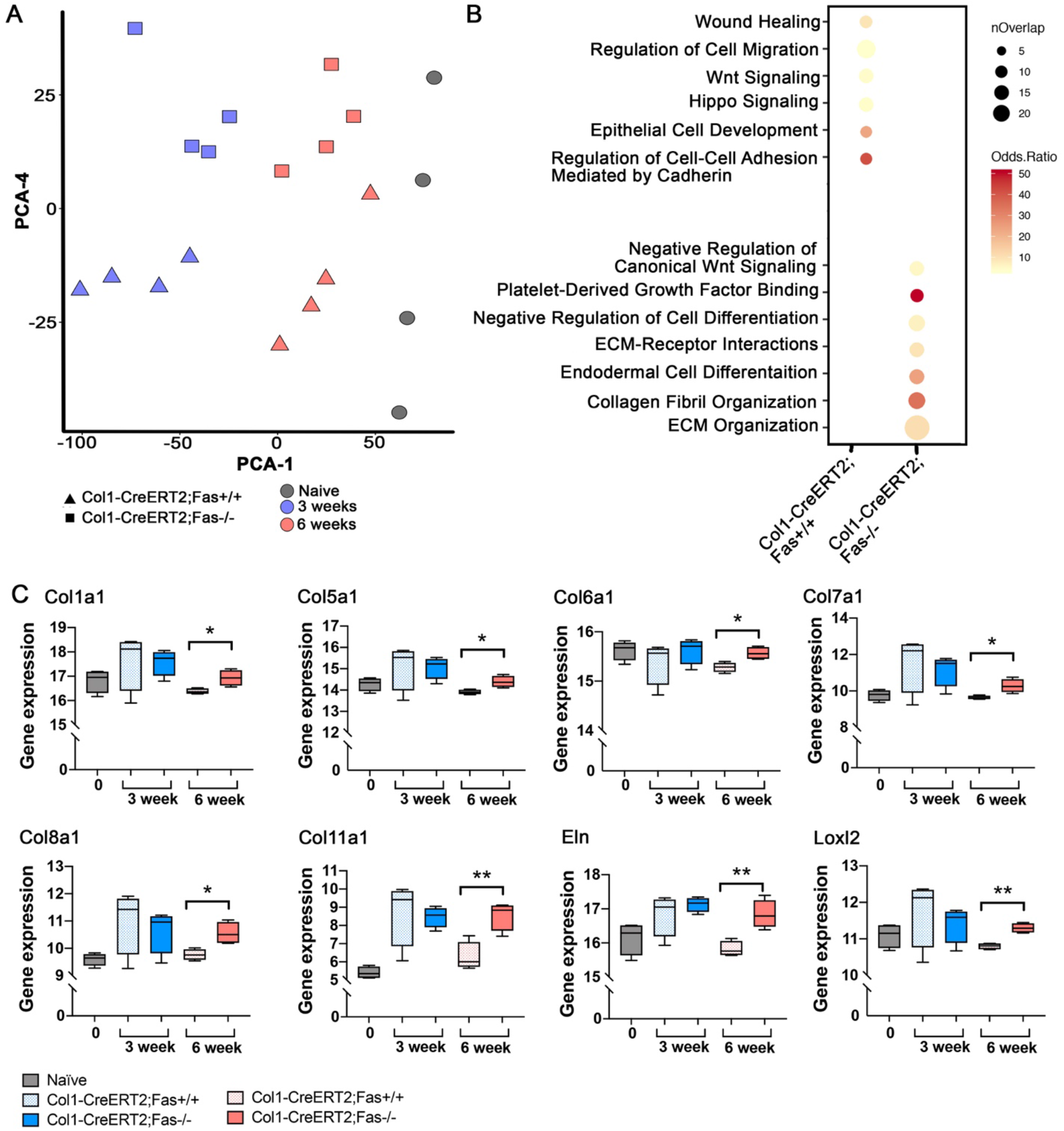
Bulk sequencing reveals a sustained pro-fibrotic signature in fibroblasts in the absence of Fas. (A) PCA plot of global transcription patterns in Col1-CreERT2;Fas^+/+^ and Col1-CreERT2;Fas^−/−^ Lin(-) fibroblasts from control and bleomycin treated mice after 3 and 6 weeks. (B) Representation of enrichment for GO pathway categories 6 weeks after bleomycin from Col1-CreERT2;Fas^+/+^ and Col1-CreERT2;Fas^−/−^ Lin(-) fibroblasts. (C) Box-and-whisker plots of genes expressed during fibrosis and in non-resolving Col1-CreERT2;Fas^−/−^ fibroblasts. (mean +/− SEM, n=4). *p<0.05, **p<0.01, Student’s t test.

To investigate heterogeneity among lung fibroblast populations, we conducted scRNA-seq on sorted Lin-cells from naïve mice and bleomycin-instilled Col1-CreERT2;Fas^−/−^ and Col1-CreERT2;Fas^+/+^ mice at 3 and 6 weeks (**Fig. 7A**). Although our scRNA-seq was conducted on purified Lin-fibroblasts, minor clusters of microvascular and lymphatic endothelial cells, alveolar epithelial cells, hematopoietic cells and erythroid cells were also identified (**Fig. S9A,B**). To focus our analysis on fibroblasts populations, we removed these non-fibroblastic cell clusters, resulting in an final dataset of cell populations highly expressing canonical fibroblast genes (PDGFRα, PDGFRβ and Acta2, **Fig. 7A and Fig S9C**). With this strategy, we identified 7 major cell groupings comprising 11 cell clusters in naïve mice and fibrotic mice. (**Fig. 7A, Fig S11,S12**).

**Figure 7.**
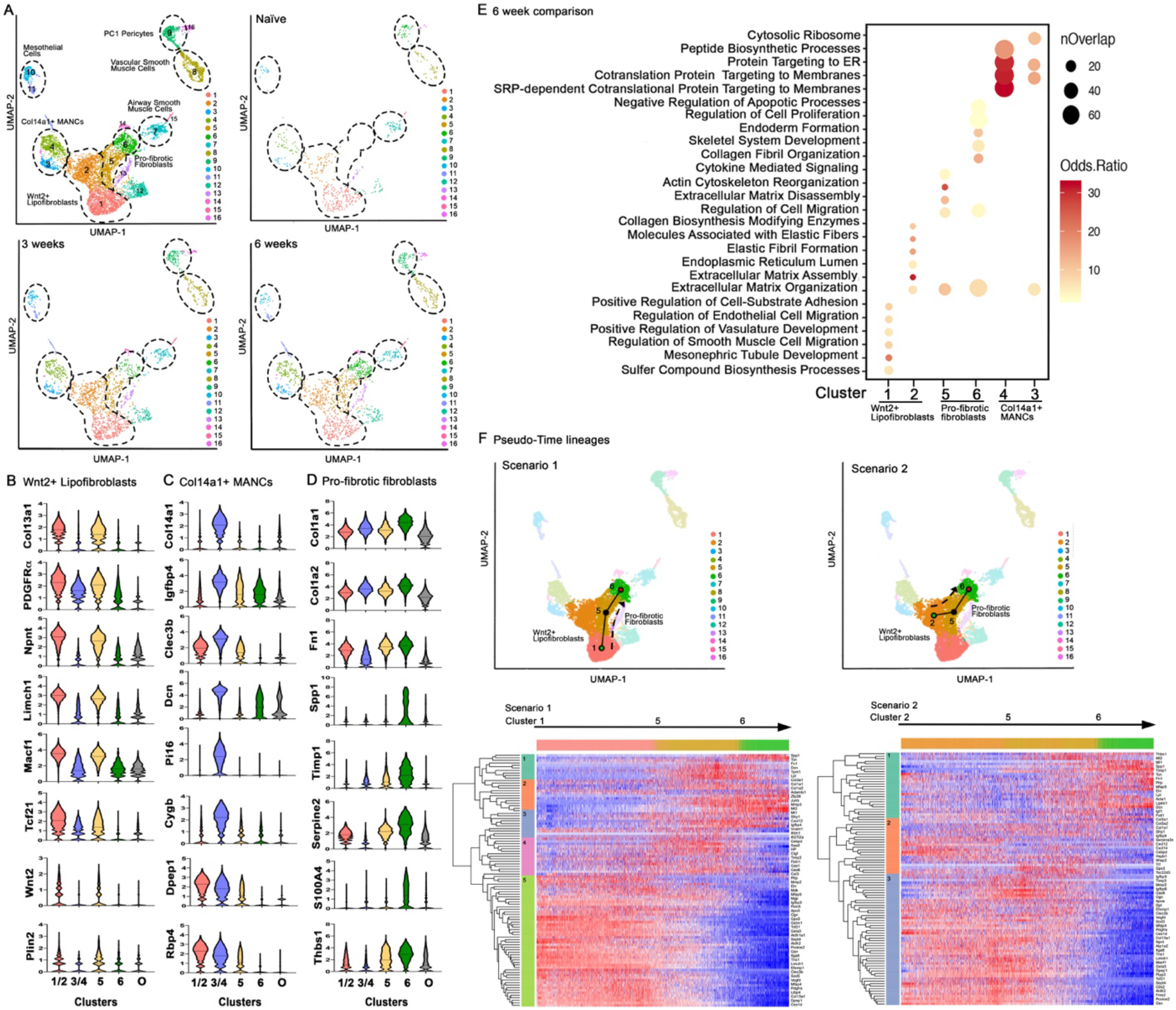
Sc-RNA sequencing reveals distinct fibroblast populations in naïve and fibrotic lungs. (A) UMAP plot of all samples and the identified population clusters and from Fas sufficient naïve mice 3 and 6 weeks after bleomycin. (B) Violin plots showing genes highly associated with Wnt2+ Lipofibroblasts (clusters 1/2). (C) Violin plots showing genes highly associated with Col14a1+ MANCs cells (clusters 3/4). (D) Violin plots showing genes highly associated with pro-fibrotic fibroblasts (clusters 5/6). O = combined expression of other clusters (3, 6, 8, 9, 11, 13, 15). (D) Representation of enrichment for GO pathway categories for clusters 1, 2, 3, 4, 5 and 6, weeks after bleomycin. (F) Pseudotime trajectory analysis and heatmap of differential expressed genes for 2 scenarios. (mean +/− SEM, n=4 pooled mice/group). ***p<0.001.

Clusters 1 and 2 bore similarity to previously described Col13a1-expressing and Wnt2+ fibroblasts (28, 29). These clusters also showed enriched expression of typical lipofibroblast genes including Plin2, Tcf21 and PDGFRα as well as other genes including Limch1 and Npnt (**Fig. 7B and Fig. S10**). We collectively define clusters 1 and 2 as a single population of “Wnt2+ lipofibroblasts”. Clusters 3 and 4 displayed similarity to both Col14a1-expressing fibroblasts and Axin2+PDGFRα+ mesenchymal alveolar niche cells (MANCs) (28, 29) and were defined by enriched expression of Col14a1, Dcn, Igfbp4, Macf1, Cygb, Clec3b and Rbp4 (**Fig. 7C and Fig S10**). Thus, we define clusters 3 and 4 as a single population of “Col14a1+ MANCs”. Cluster 7 was identified as airway smooth muscle cells and/or PC2 pericyte-like cells based on enriched expression of Hhip, Must1, Aspn, Enpp2, Lum and Acta2, (**Fig. 7A and Fig S11D**); while cluster 8 was identified as vascular smooth muscle cells and Axin+ myogenic progenitor cells based on enriched expression of Actc1, Acta2, Myh11, CNN1, Itih4, Lmod1 and Lgr6 (**Fig. 7A and Fig S11B**). Cluster 9 was identified as PC1 pericyte-like cells which displayed abundant expression of Pdzd2, Cox4i2, Adcy8 and Higd1b along with the highest expression of PDGFRβ among all clusters (**Fig. S10B and S11C**) (28, 44, 45); while clusters 10 and 11 were identified as single group of mesothelial cells based on enriched expression of Upk3b, Lrrn4, Msln, Gpm6a and Rspo1 (**Fig. 7A and Fig. S12E**). An additional, novel population comprising clusters 5 and 6 was identified in the lungs of bleomycin-instilled mice (**Fig. 7A**). Cluster 5 exhibited increased expression of archetypal profibrotic genes including Col1a1, Col1a2, and Tbsp1, whereas cluster 6 was defined by enrichment with additional and distinct pro-fibrotic genes, including Spp1, Fn1, Cthrc1, and Thbs1. In addition, cluster 6 exhibited the highest expression of PDGFRα as well as substantially increased Acta2 expression among fibroblast clusters (**Fig. 7D**). Based on these characteristics, we define clusters 5 and 6 as “pro-fibrotic fibroblasts”.

Next, we compared the gene expression patterns among the Wnt2+ lipofibroblasts, Col14a1+ MANCs and pro-fibrotic fibroblast subsets in bleomycin-instilled Col1-CreERT2;Fas^−/−^ and Col1-CreERT2;Fas^+/+^ mice at 3 and 6 weeks. To verify efficient deletion of Fas in Col1-CreERT2;Fas^−/−^ mice, we compared the gene expression pattern of the deleted exon 9 between Col1-CreERT2;Fas^−/−^ and Col1-CreERT2;Fas^+/+^ mice at 3 and 6 weeks. Expression of exon 9 was significantly reduced (p<0.001) and the number of cells expressing exon 9 was decreased 4.8-fold (from 39% to 8%) at 3 weeks and 2.3-fold (from 43% to 19%) at 6 weeks in Fas-deficient cells (**Fig. S13**). Exon 9 was deleted similarly in all clusters in Col1-CreERT2;Fas^−/−^ mice (not shown). Similar to the differential expression analysis seen by bulk RNA-seq, Fas-deficiency in fibroblasts had little impact on the induction of genes related to pro-fibrotic pathways at 3 weeks in any of the fibroblast subsets, consistent with our data showing that fibroblast Fas-deficiency had no effect on the development of fibrosis. By contrast, comparison of specific fibroblast populations between Col1-CreERT2;Fas^−/−^ and Col1-CreERT2;Fas^+/+^ mice at 6 weeks revealed fibrosis-associated DEGs in the pro-fibrotic fibroblasts (clusters 5 and 6; 40 genes) and the Wnt2+ lipofibroblasts (clusters 1 and 2; 95 genes), but not in the Col14a1+ MANCs (clusters 3 and 4). Within the pro-fibrotic fibroblasts (clusters 5 and 6) of Col1-CreERT2;Fas^−/−^ mice at 6 weeks, gene ontology pathways associated with extracellular matrix organization (p=1.916E-29), regulation of cell migration and actin cytoskeleton reorganization (p=5.711E-07) and negative regulation of apoptotic processes (p=3.599E-06) were also highly enriched, indicating that pro-fibrotic signatures persisted in this Fas-deficient pro-fibrotic subset (**Fig. 7E and Table S2**). Additionally, the upregulated DEGs from the Wnt2+ lipofibroblasts were enriched for gene ontology pathways associated with elastic fibril formation (p=0.0072), collagen biosynthesis modifying enzymes (p=0.0335), and containing other extracellular matrix assembly genes (Col28A1, Adamts2, Col13A1, Col4A5, Cst3, Rcn1, Sparcl1, Igfbp7, Gas6, Vcam1, Mmp2, Mmp3, Loxl1, Fbln5). Gene ontology pathways associated with peptide biosynthesis processes, and targeting of proteins to the ER and membranes were enriched in Col14a1+ MANCs fibroblasts (**Fig. 7E and Table S2**). Thus, both the pro-fibrotic fibroblasts and the Wnt2+ lipofibroblasts retained pro-fibrotic gene expression patterns in the absence of Fas-signaling.

Lastly, to provide insight into the potential progenitors and lineages leading to the pro-fibrotic fibroblast populations (clusters 5 and 6) that emerge with injury and fibrosis, we performed in silico trajectory analyses of our single cell data. We included the Wnt2+ lipofibroblasts population (clusters 1 and 2) in this analysis as potential progenitors given their related expression profile, exemplified by the connected nature of clusters 1, 2, 5, and 6 in UMAP space. Further supporting cluster 1 as a progenitor for clusters 5 and 6, we found the frequency of this cluster decreased from 30% in naïve animals to 20% and 14% as fibrosis developed at 3 and 6 weeks respectively. Pseudotime trajectory analysis inferred two major trajectories: (Lineage 1): proceeding from cluster 1 through cluster 5 to cluster 6, and (Lineage 2): proceeding from cluster 2 through cluster 5 to cluster 6 (**Fig. 7F**). Through clustering of pseudotime-associated genes, we discretized the trajectories into early, mid, and late phases. The pseudotime-associated genes for the early and mid phases of Lineage 1 exhibited enrichment for pro-fibrotic pathways including lung fibrosis (p=0.036), extracellular matrix-receptor interactions (p=8.18E-08), focal adhesion formation (p=3.4E-06), TGF-β signaling pathways (p=0.05327) and matrix metalloproteinases (p=0.01727) (**Fig. 7F, Table S3**). Similar pathways were enriched during the early and mid phases of the lineage 2, but also included genes involved in glutathione metabolism (Gpx3) and VEGF signaling (Hspb1). Late trajectory genes included matrix metalloproteinase genes (Mmp3, Timp3) and genes involved in IGF1 signaling (Igfbp3, Igfbp6) (**Fig. 7F Table S4**). Thus, pseudotime analysis suggests that the pro-fibrotic fibroblasts (clusters 5 and 6) develop from the Wnt+ lipofibroblasts (clusters 1 and 2) in naïve lungs. Taking into consideration both the bulk and scRNA-seq data, our findings suggest that once each fibroblast cluster has undergone its unique pro-fibrotic programming response, the pro-fibrotic gene expression profile remains largely unchanged as long as fibrosis persists.

## DISCUSSION

Substantial progress has been made in understanding the mechanisms of alveolar epithelial injury and its role in driving fibroblast recruitment, proliferation and excessive ECM deposition in the distal lung. However, the mechanisms that distinguish pro-fibrotic fibroblast persistence and ECM accumulation in the context of progressive pulmonary fibrosis seen in IPF from the fibrosis resolution and lung regeneration that occurs in the setting of recovery from ARDS, remain unclear. Here, we show that Fas deletion in pro-fibrotic fibroblasts inhibits fibroblast apoptosis, impedes homeostatic fibrosis resolution, maintains pro-fibrotic transcriptomic fibroblast gene expression programing and permits fibroblast persistence and enduring pulmonary fibrosis. Taken together, these findings suggest that Fas signaling plays a fundamentally important, physiologic role in the elimination of pro-fibrotic lung fibroblasts during fibrosis resolution *in vivo*. Furthermore, our results suggest that impaired Fas signaling e.g. by Fas down-regulation or by increased expression of anti-apoptotic proteins that inhibit pro-apoptotic Fas signaling (10, 16–19), may lead to critical deviations in lung fibrosis outcome from resolution to persistence.

Two approaches were used to delete Fas in fibroblasts. Dermo1 is developmentally expressed in lung mesenchyme (25). Thus, fibroblasts in Dermo1-Cre;Fas^−/−^ mice are Fas-deficient at birth (46–48). We also conditionally deleted Fas in fibroblasts during bleomycin-induced fibrosis development in Col1-CreERT2;Fas^−/−^ mice. With both approaches, bleomycin robustly induced pulmonary fibrosis 2-4 weeks. However, whereas Fas-sufficient mice underwent the homeostatic fibrosis resolution by 6-9 weeks (8, 9, 23, 49), fibroblast Fas deficiency in both Dermo1-Cre;Fas^−/−^ and Col1-CreERT2;Fas^−/−^ mice inhibited homeostatic fibrosis resolution and led to persistent pulmonary fibrosis for at least 9 weeks.

Fibroblast Fas deficiency profoundly inhibited both the apoptosis and elimination of lung fibroblasts during homeostatic fibrosis resolution. Using a novel flow cytometry strategy for pulmonary fibroblasts, based on previous studies describing CD90+, CD90- and CD26+ fibroblasts in lung and skin (34, 50), we found that among Fas-deficient lung fibroblasts, CD26+ and CD90-CD26-fibroblasts remained the most persistent and elevated subsets at 6 and 9 weeks. CD90+ fibroblasts underwent a modest early expansion, but also remained significantly elevated at 6 and 9 weeks in the absence of Fas signaling. Furthermore, whereas apoptotic S100A4^+^ and αSMA^+^ fibroblasts were detected in the lungs of bleomycin-instilled Dermo1-Cre;Fas^+/+^ and Col1-CreERT2;Fas^+/+^ mice at 3 and 4.5 weeks, they were not detected in Dermo1-Cre;Fas^−/−^ and Col1-CreERT2;Fas^−/−^ mice suggesting that Fas-signaling plays an essential role in fibroblast apoptosis during homeostatic fibrosis resolution. The lungs of naïve Dermo1-Cre;Fas^−/−^ mice were found to have basally elevated numbers of Lin-PDGFRα+ cells compared to naïve Dermo1-Cre;Fas^+/+^ mice, with the majority being CD26+. These findings suggest that Fas signaling may be of greater relevance to basal turnover of CD26+ fibroblasts compared to CD90+ and CD90-CD26-fibroblasts. However, we did not observe spontaneous lung or skin fibrosis in these mice or in aged (i.e. >1-year old) Dermo1-Cre;Fas^−/−^ mice (not shown). We also noted that fibroblast Fas-deficiency in Dermo1-Cre;Fas^−/−^ and Col1-CreERT2;Fas^−/−^ mice had no effect on the initial bleomycin-induced fibroblast expansion at 1.5 and 3 weeks. Previous *in vitro* studies have suggested that Fas-signaling during this period contributes to pro-fibrotic fibroblast proliferation via cFLIPL and TRAF2-induced NF-κB activation (19). Our use of genetic approaches to delete Fas in fibroblasts clearly challenge this notion. Taken together, our results suggest that Fas signaling plays a necessary role in fibroblast apoptosis and elimination during homeostatic fibrosis resolution following bleomycin-induced lung injury and likely contributes to fibroblast turnover in naïve mice.

Much remains to be understood about the functions of the increasingly heterogeneous lung fibroblast subsets. We initially studied how Fas-signaling affects CD90+, CD26+ and CD90-CD26-fibroblasts. CD90 is expressed by Tcf21-expressing lipofibroblasts and Col13a1 interstitial matrix-producing fibroblasts (51). In the context of bleomycin-induced pulmonary fibrosis, CD90-deficient mice phenocopy the impaired homeostatic fibrosis resolution described herein, and lung fibroblasts from CD90-deficent mice are resistant to Fas-induced apoptosis (9, 52). CD26 has been reported to be expressed on dermal fibroblasts that synthesize and deposit ECM during scar formation (34) and we have shown it to identify a CD90-fibroblast subset. We have also shown that CD90+, CD26+ and CD90-CD26-fibroblasts from bleomycin-instilled mice exhibit increased Col1a1-promoter activity, peaking at 3 weeks in Col1-GFP reporter mice. However, whereas GFP had returned to baseline level by 6 weeks in Fas-sufficient fibroblasts from Col1-CreERT2;Fas^+/+^;Col1-GFP mice, GFP expression remained elevated in Fas-deficient CD90+, CD26+ and CD90-CD26-fibroblasts in Col1-CreERT2;Fas^−/−^;Col1-GFP mice. Similarly, αSMA promoter activity remained elevated in CD90+ fibroblasts in bleomycin-instilled Fas-deficient Col1-CreERT2;Fas^−/−^;αSMA-RFP reporter mice 6 weeks compared to CD90+ fibroblasts from Fas-sufficient Col1-CreERT2;Fas^+/+^;αSMA-RFP reporter mice. Thus, loss of Fas-signaling in fibroblasts not only prevents fibroblast apoptosis, it prolongs functional gene expression from pro-fibrotic Col1a1 and αSMA promoters in surviving cells.

Bulk and scRNA-seq provided further insight into the transcriptional profiles and, by inference, the functions of the Fas-deficient fibroblasts that remained in persistently fibrotic lungs. Analysis of bulk RNA-seq data from Fas-sufficient and Fas-deficient fibroblasts showed marked similarities in pro-fibrotic gene expression and pathway enrichment at 3 weeks compared to naïve mice. scRNA-seq provided additional insight. We identified two major fibroblast subsets in the lungs of naïve mice. “Wnt2+ lipofibroblasts” (clusters 1 and 2) displayed similarity to Wnt2+ (29) and Col13a1 fibroblasts (28), but were also enriched with transcripts encoding lipofibroblast genes including Plin2 (28, 51). “Col14a1+ MANCs” were similar to Axin+PDGFRα+ and Col14a1+ fibroblasts (28, 29). At the peak of fibrosis, Wnt2+ lipofibroblasts from both Fas-sufficient and Fas-deficient mice displayed increased expression of typical pro-fibrotic genes. In contrast, gene expression profiles in Col14a1+ MANCs from bleomycin-instilled mice did not display a gene expression profile associated with the development or maintenance of fibrosis and very few DEGs were found in comparision with lung fibroblasts from naïve mice, as previously reported (29). We also identified a third, novel, pro-fibrotic fibroblast population (clusters 5 and 6) that was present in the lungs of both Fas-sufficient and Fas-deficient bleomycin-instilled mice, but was absent from the lungs of naïve mice. Our data supports the conclusion that these populations arise from the Wn2t+ lipofibroblasts (clusters 1 and 2) present in naïve lungs. *In silico* trajectory analysis provided two possible differentiation paths initating with either cluster 1 or 2, and passing through cluster 5 to generate the pro-fibrotic cluster 6 cells.

Analysis of the lung gene expression profiles in fibroblasts isolated from bleomycin-instilled Col1-CreERT2;Fas^+/+^ mice at 6 weeks showed enrichment in pathways associated with regulation of cell migration and motility, wound healing, Wnt signaling and epithelial cell development, consistent with homeostatic fibrosis resolution. However, the Wnt2+ lipofibroblasts (clusters 1 and 2) and the pro-fibrotic fibroblasts (clusters 5 and 6) isolated from fibrotic bleomycin-instilled Col1-CreERT2;Fas^−/−^ mice continued to express pro-fibrotic gene expression patterns and pathways at 6 weeks, indicating that their pro-fibrotic programming had been preserved in the absence of Fas-signaling.

Taken together, our data suggest that following initial pro-fibrotic programming of Wnt2+ lipofibroblasts during the development of fibrosis, these cells and the novel pro-fibrotic fibroblasts that are derived from them maintain their pro-fibrotic transcriptomic programming. Therefore, autonomous Fas-induced apoptosis and clearance of pro-fibrotic fibroblasts during fibrosis resolution plays a necessary role in homeostatic fibrosis resolution, whereas impaired Fas-signaling is sufficient to maintain both their presence in persistently fibrotic lungs and their pro-fibrotic programming. We speculate that loss of the ability of fibroblasts to undergo Fas-induced apoptosis represents a decisive checkpoint at which the beneficial, resolving fibrotic response seen in ARDS survivors, might be diverted to the persistent and potentially more harmful progressive fibrosis seen in IPF patients.

## METHODS

### Mouse strains

To enact lineage specific deletion of Fas, we bred mice that expressed endogenous Cre recombinase driven by the *Dermo1 (Twist2)* promoter (008712 - B6.129X1-*Twist2^tm1.1(cre)Dor^*/J; Jackson Laboratories) or a tamoxifen-inducible Cre recombinase driven by the Col1a1 promoter (016241 - B6.Cg-Tg(Col1a1-cre/ERT2)1Crm/J; Jackson Laboratories, Bar Harbor, ME) to mice with floxed Fas alleles (exon 9) (007895-C57BL/6-*Fas^tm1Cgn^*/J; Jackson Laboratories). The final genotypes of the mice were *Dermo1-Cre;Fas^fl/fl^* and *Col1a1-Cre-ERT2;Fas^fl/fl^* respectively. To track real-time expression of Col1a1 and αSMA, we used Col1a1-GFP and αSMA-RFP mice (a gift from David Brenner, (39)). The Col1a1-GFP and αSMA-RFP mice were bred to Col1a1-Cre-ERT2;Fas^fl/fl^ to allow for Fas deletion and florescence expression of Col1a1 or αSMA. For all mice bred to the Col1a1-ERT2 promoter either corn oil or tamoxifen (0.25 mg/g body weight in corn oil, Sigma Aldrich, St. Louis, MO) was given by intraperitoneal injection starting at day 4 after bleomycin. Mice received 6 injections between day 4 and 21 (**Fig. 1F**). Deletion of Fas after Cre recombination following tamoxifen was confirmed by RT-PCR (**Supplementary Fig. 2**, Cre Fwd – GAGTGAACGAACCTGGTCGAAATCAGTGCG; Cre Rev – GCATTACCGGTCGATGCAACGAGTGATGAG; Fas Del Fwd – GTCCTCTATTATCCTCATCATGAG; Fas Del Rev – GGCTTTGGAAAGGAATTTCCTCCTAAGAGG; Fas LoxP Fwd – CCTTCCATTGATGGACAGTTC; Fas LoxP Rev – TTAAAAGGCTTTGGAAAGGAA). All Animal studies were approved by the National Jewish Health Institutional Animal Care and Use Committee.

### Primary Fibroblasts

Primary lung fibroblasts were isolated from healthy murine lungs as previously described (16, 53). They were maintained in 10% DMEM and grown on plastic and used in experiments between passages 3-8.

### Assessment of fibrotic lung disease

Pulmonary fibrosis was initiated by the intratracheal instillation of 50 μl of bleomycin (1.5U/kg, Amneal Biosciences, Bridgewater, NJ) to anesthetized mice, as previously described (23). Fibrosis was assessed by lung measurements of collagen in the upper right lobe (hydroxyproline). Briefly, lungs were homogenized in PBS and hydrolyzed overnight at 120°C using 12M HCl. Assessment of hydroxyproline was determined by the absorbance at 500nm on a microplate reader (Epoch2 BioTek Instruments, Winooski, Vermont). Histology was evaluated by Hematoxylin and Eosin (H&E) staining and Picrosirius red staining (PSR) of sections from the left lung as previously described (23). Images were taken on an upright Olympus BX51.

### Flow cytometry and cell sorting

Single cell suspensions were obtained from perfused, enzymatically dispersed lungs. Briefly, a digestion mixture of collagenase (450 U/ml), Dispase (5U/ml) and Elastase (4U/ml) was instilled into the right lung and incubated at 37°C for 25 min. Lungs were chopped and tissue pieces washed in 10% FBS containing DMEM followed by a secondary digestion in 0.1% Trypsin-EDTA with 0.33 U/ml DNaseI for 20 min at 37°C. Dissociated tissue was washed in DMEM containing 10% FBS and single cell suspensions were filtered prior to staining (36). Cells were stained with fluorescently-tagged monoclonal antibodies against CD45, CD31, CD326/EpCAM, CD90, CD26, and PDGFRα/CD140a (ThermoFisher, Waltham, MA) at a 1:200 dilution. Cell analysis data were acquired with the LSRFortessa (BD Biosciences, San Jose, CA) and analyzed with FlowJo 2 software (Tree Star, Ashland, OR).

Prior to cell sorting, single cell suspensions were enriched for linage negative cells by incubating with CD45, CD31 and CD326 MicroBeads and purified off of LS columns per manufacturer’s instructions (Miltenyi Biotech, Bergisch Gladbach, Germany). Single cells were sorted on the FACSAria Fusion (BD Biosciences) as doublet excluded, DAPI negative, CD45, CD31, and CD326 negative prior to single cell and bulk RNA sequencing of the lineage negative population.

### Immunofluorescent staining

Formalin fixed, paraffin embedded sections, were deparafinized and rehydrated followed by antigen retrieval using citrate buffer. Non-specific binding was reduced by incubation in 10% blocking serum and mouse IgG serum (Vector Laboratories, Burlingame, CA). Primany anti-rabbit S100-A4/FSP1 antibody (Milliopore Sigma, Burlington, MA) was used at a 1:500 dilution and primary anti-mouse αSMA antibody (SigmaAldrich) was used at a 1:1000 dilution overnight at 4°C followed by incubation with fluorescently tagged goat anti mouse-A647 and donkey anti-rabbit-A555 secondary antibodies at 1:100 dilution (Invitrogen, Carlsbad, CA). TUNEL staining to detect apoptotic cells was completed prior to antibody stainig per manufactuors instructions (Promega, Madison, WI). TUNEL positive cells were counted and averaged from 10 images from each animal. Images were captued in fibrotic ares using FSP1 positive staining as an indicator of fibroblast accumlation. TUNEL positive cells located in the airway epithelium were excluded from the analysis. Frozen 4% PFA fixed OCT (Fisher Scientific, Hampton, NH) sections were placed on slides, dried and OCT was removed by washing in PBS. Slides were mounted with Fluoroshield Mounting Media containing DAPI (Vector Laboratories). Images were acquired on a Zeiss Axioplan 2 epi-fluorescence microscope and analyzed with Axiovision software (Zeiss, Jena, Germany).

### Bulk RNA sequencing

Lineage negative cells were obtained by FACs sorting. At least 300,000 total cells were collected per population. Purified cells were pelleted, lysed and RNA was precipitated using TRIzol (ThermoFisher Scientific) followed by purification of the aqueous layer using a RNeasy Micro Kit (Qiagen Sciences, Valencia, CA). A modified Kapa Biosystems (Wilmington, MA, USA) KAPA Stranded mRNA-Seq kit for whole transcriptome libraries was used to primarily target all polyA RNA. After mRNA (poly-A) isolation, 1^st^ and 2^nd^ strand cDNA synthesis, adaptor ligation, amplification, and bead templating occurred. Once validated, the libraries were sequenced as barcoded-pooled samples and processed for next-generation sequencing (NovaSeq 6000 - Illumina platform). The data set has been deposited in the National Center for Biotechnology Information/Gene Expression Omnibus under accession number GEO:XXXX

### Single cell RNA sequencing

All single-cell capture and library preparation was performed at the University of Colorado Cancer Center Microarray and Genomics Core. Cells were loaded onto a 10x Genomics microfluidics chip and encapsulated with barcoded oligo-dT-containing gel beads using the 10x Genomics Chromium controller and single-cell libraries constructed per the manufacturer’s instructions. Samples were sequenced on a NovaSeq 6000 Illumina platform instrument (Illumina, San Diego, CA). An average of 2741 cells per sample were analyzed with an average post normalized mean reads/cell of 312,995. The data set has been deposited in the National Center for Biotechnology Information/Gene Expression Omnibus under accession numbers GEO:XXXX

### Statistical Analysis

Data are presented as the mean ± SEM and analyzed using GraphPad Prism software Version 8 (GraphPad, San Diego, CA). Differences between conditions at specific time points were examined using Student’s unpaired t-test. One-way ANOVA with Newman-Keuls post-hoc analysis was used to compare results from more than two groups with p<0.05 considered to be significant. Specifics about the replicates used where n = individual animal replicates are available in the Figure legends.

### Bulk RNA sequencing analysis

To improve downstream mapping quality, raw sequencing reads were trimmed using skewer with parameters (end-quality=15, mean-quality=25, min=30). Trimmed reads were aligned to the mouse reference genome GRCm38 using Hisat2. Gene quantification was performed with htseq-count using GRCm38 ensembl v84 GTF. Differential expression analysis between groups was conducted with R package DESeq2. Pathway analsys was conducted with Enricher (54, 55).

### Single cell clustering and trajectory analysis

Initial processing of 10X scRNA-seq data, including cell demultiplexing, alignment to the mouse genome GRCm38, and UMI-based quantification was performed with Cell Ranger (version 3.0). To ensure that high quality cells were used for downstream analysis, we removed cells with fewer than 200 genes detected or cells with greater than 20% mitochondrial reads. Additionally, to remove possible doublets, we remove cells with higher than 7500 genes detected. For gene filtering, we remove lowly expressed genes (detected in fewer than 4 cells). Using the above filtering, we have a dataset consisted of 12,043 cells and 37,887 genes. After initial clustering and visualization, 4,215 cells were removed that we characterized as non-fibroblasts (**Supplemental Fig. 9**) which left us with 7,828 mesenchymal cells for downstream analysis.

Prior to clustering, we performed normalization using SCTransform and integration of datasets from five samples (naïve *Col1a1-Cre-ERT2;Fas^fl/fl^*, Col1-CreERT2;Fas^−/−^ and Col1-CreERT2;Fas^+/+^ 3 weeks after bleomycin and Col1-CreERT2;Fas^−/−^ and Col1-CreERT2;Fas^+/+^ 6 weeks after bleomycin) using mutual nearest neighbor (MNN) based approach. Clustering analysis was performed on the top 20 PCs using a shared nearest neighbor (SNN) based SLM algorithm. Visualization of the single cell expression profiles into a two-dimensional map was computed using UMAP (Uniform Manifold Approximation and Projection) technique. Differential expression analysis was conducted using FindMarkers function with default options. All the analyses mentioned above is carried out with R Seurat package version 3.5.1. Pathway analsys was conducted with Enricher (54, 55).

Using the previously computed UMAP components, trajectory analysis was performed using Slingshot that build lineages of cells that link cell clusters together by fitting a minimum spanning tree (MST) onto the selected clusters followed by the application of simultaneous principal curves to create trajectories. Clusters 1, 2, 5, and 6 were included in the trajectory analysis. Additional constraints were added by imposing cluster 1 and cluster 2 as the starting clusters and cluster 6 as the end cluster.

---

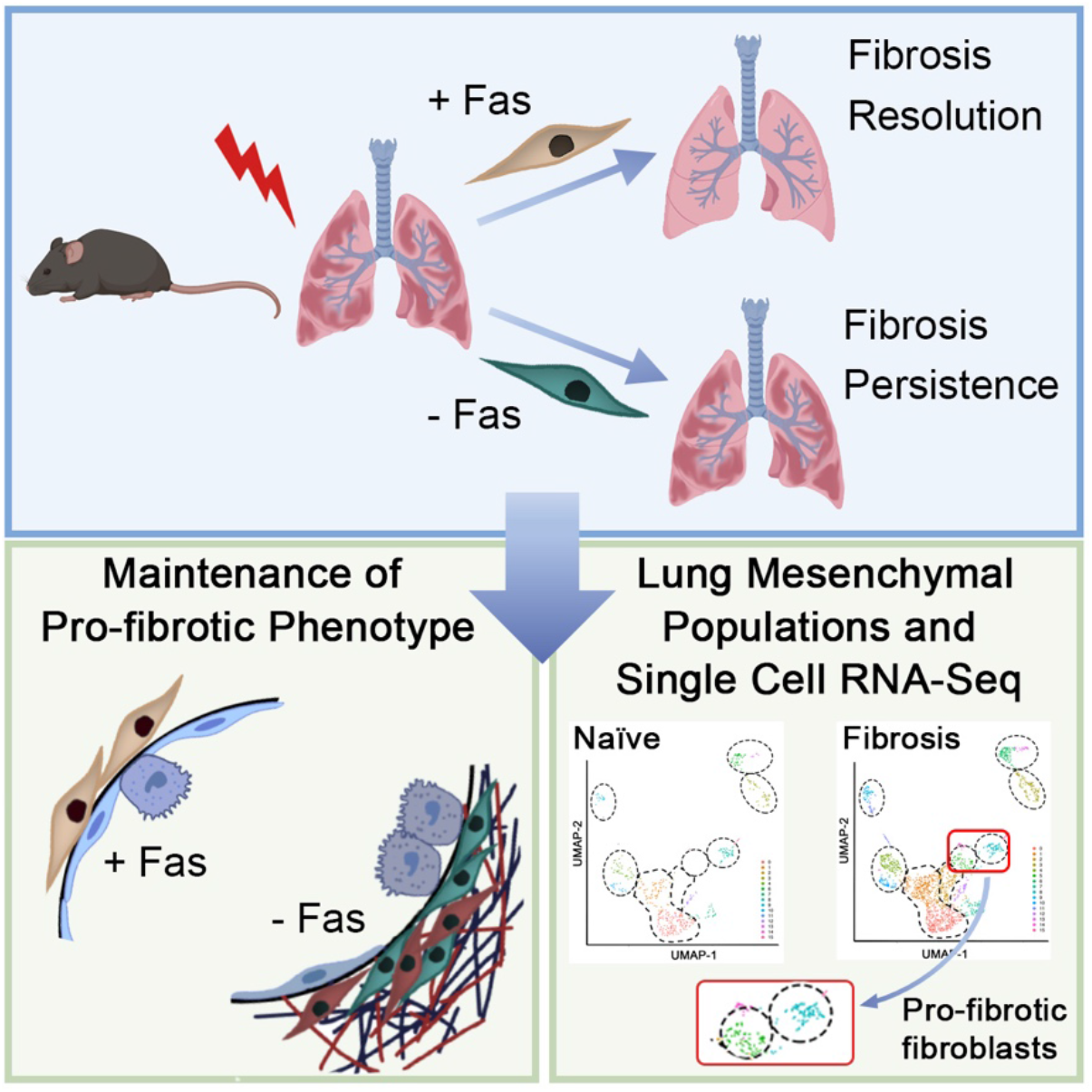
Graphical Abstract.

**Supplementary Figure 1.**
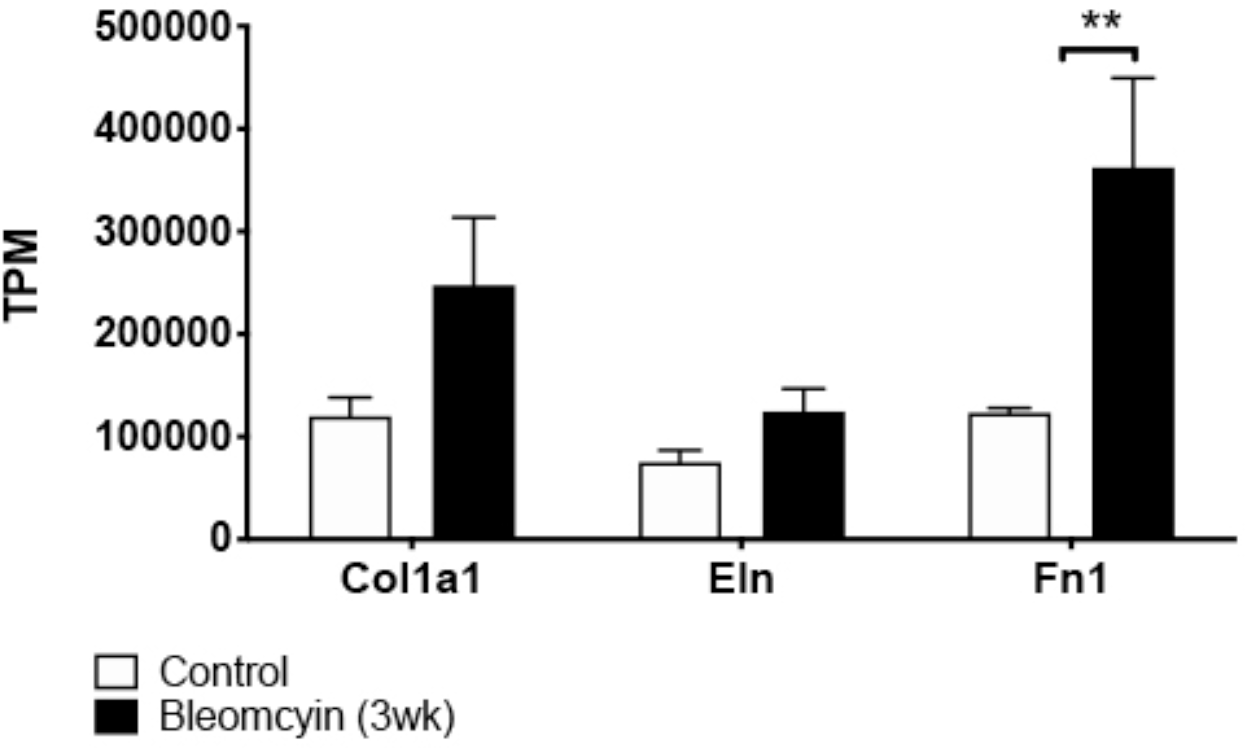
Bleomycin-induced fibrosis increases pro-fibrotic gene expression in fibroblasts. Gene expression in primary fibroblasts isolated and grown in culture from lungs of control and fibrotic lungs. (mean +/− SEM, n=3) **p<0.01, Student’s t test.

**Supplementary Figure 2.**
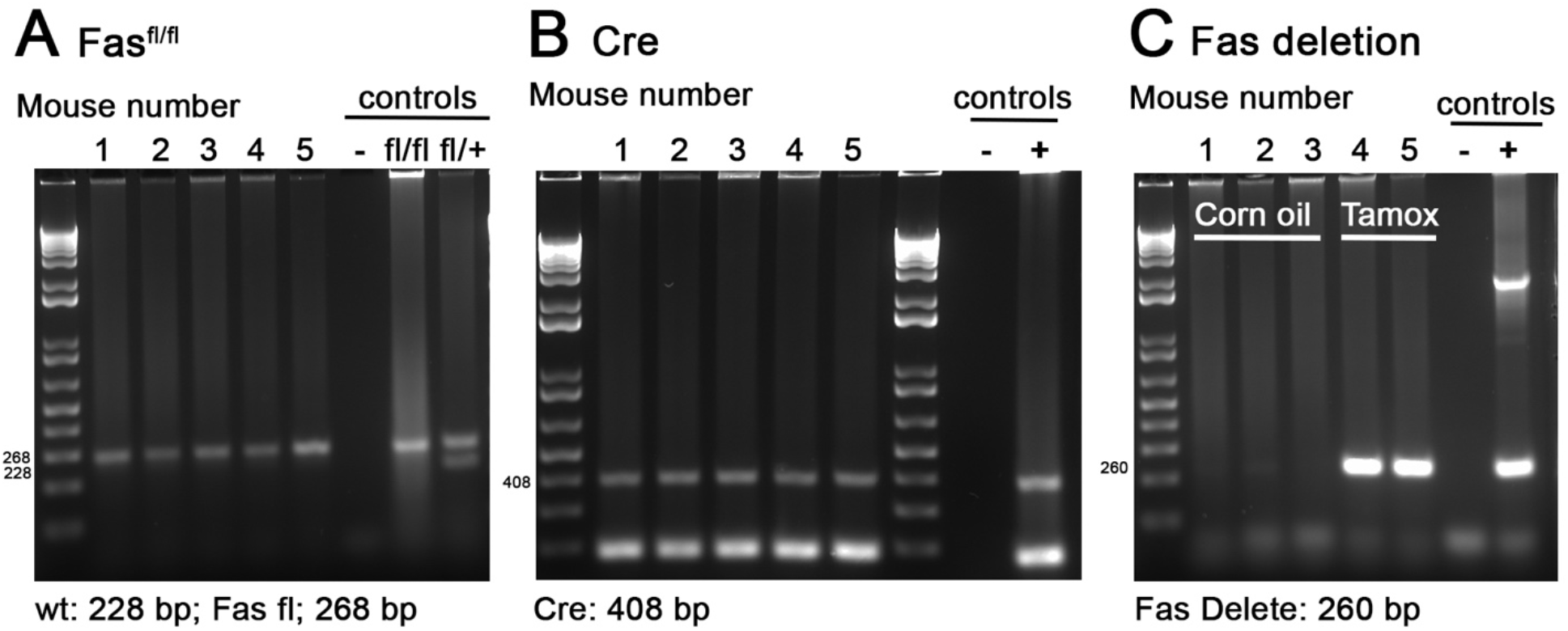
Deletion of Fas under the Col1a1 promoter after tamoxifen. (A) PCR genotyping of loxp sights around exon 9 of Fas. (B) PCR genotyping of Cre. (C) PCR genotyping of Fas deletion band after *in vivo* tamoxifen treatment.

**Supplementary Figure 3.**
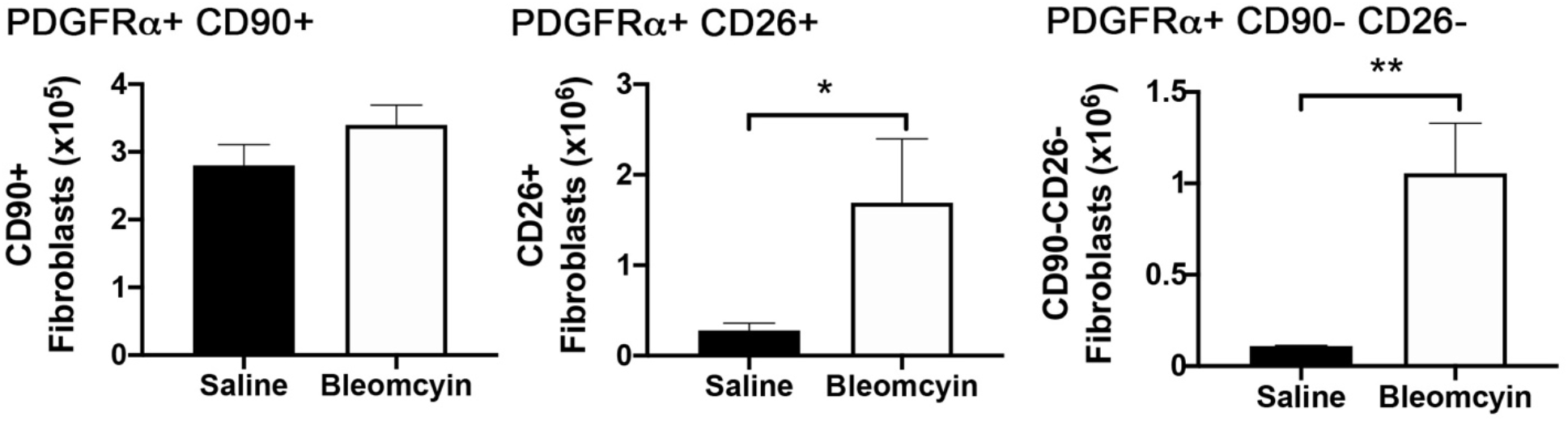
Fibroblast subpopulations in fibrotic wild type mice. Quantitation of Lin-PDGFRα+ fibroblast sub-populations: CD90+, CD26+ and CD90-CD26-by flow cytometry. (mean +/− SEM, n=3). *p<0.05, **p<0.01.

**Supplementary Figure 4.**
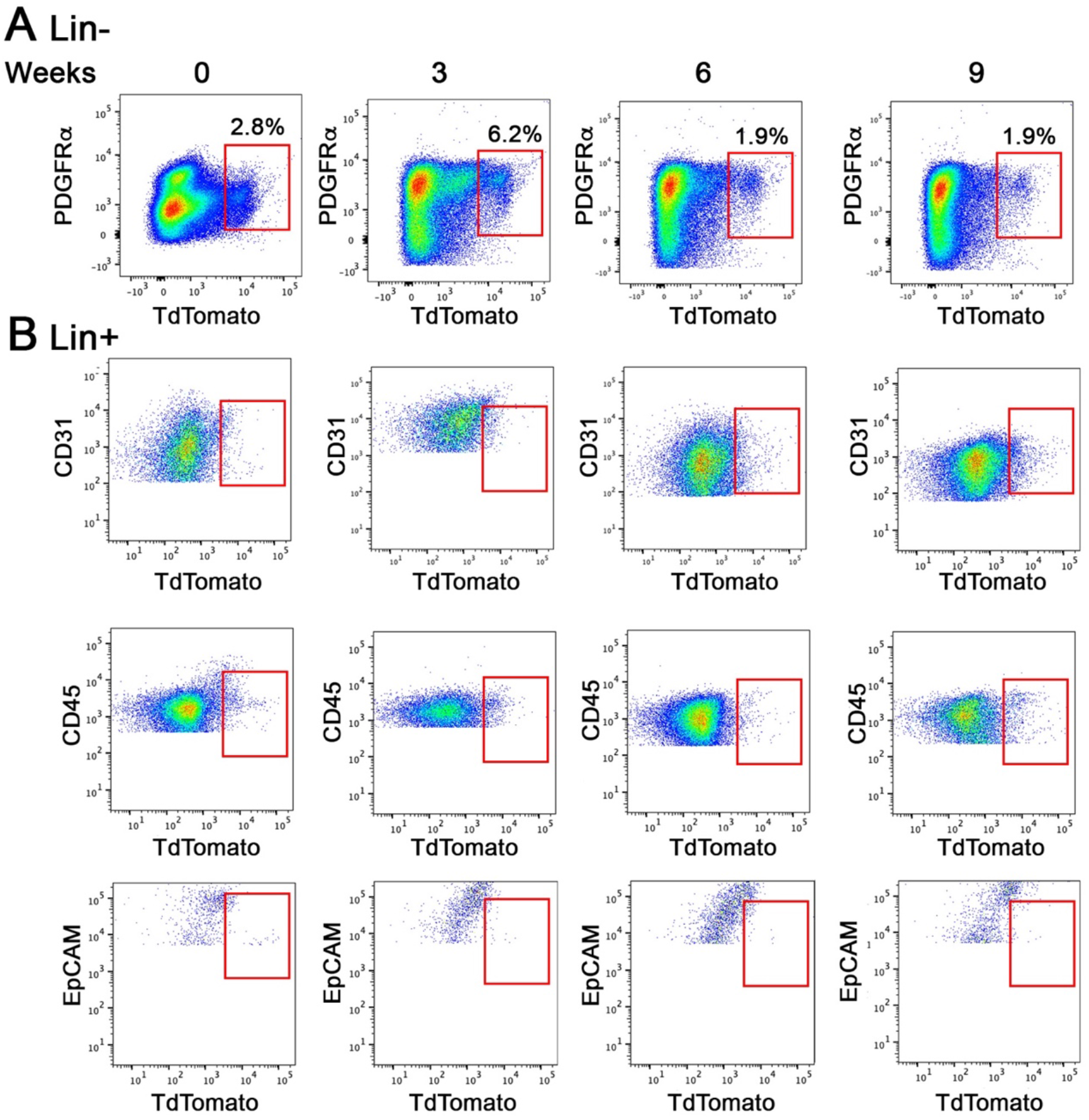
Flow cytometry gating strategy for Lineage positive cells. (A) Representative gating strategy to identify lineage positive CD31, CD45 and EpCAM cells. (B) Representative flow plots of GFP expression in lineage positive cells over time after bleomycin in Col-GFP mice. (C) Representative flow plots of RFP expression in lineage positive cells over time after bleomycin in αSMA-RFP mice.

**Supplementary Figure 5.**
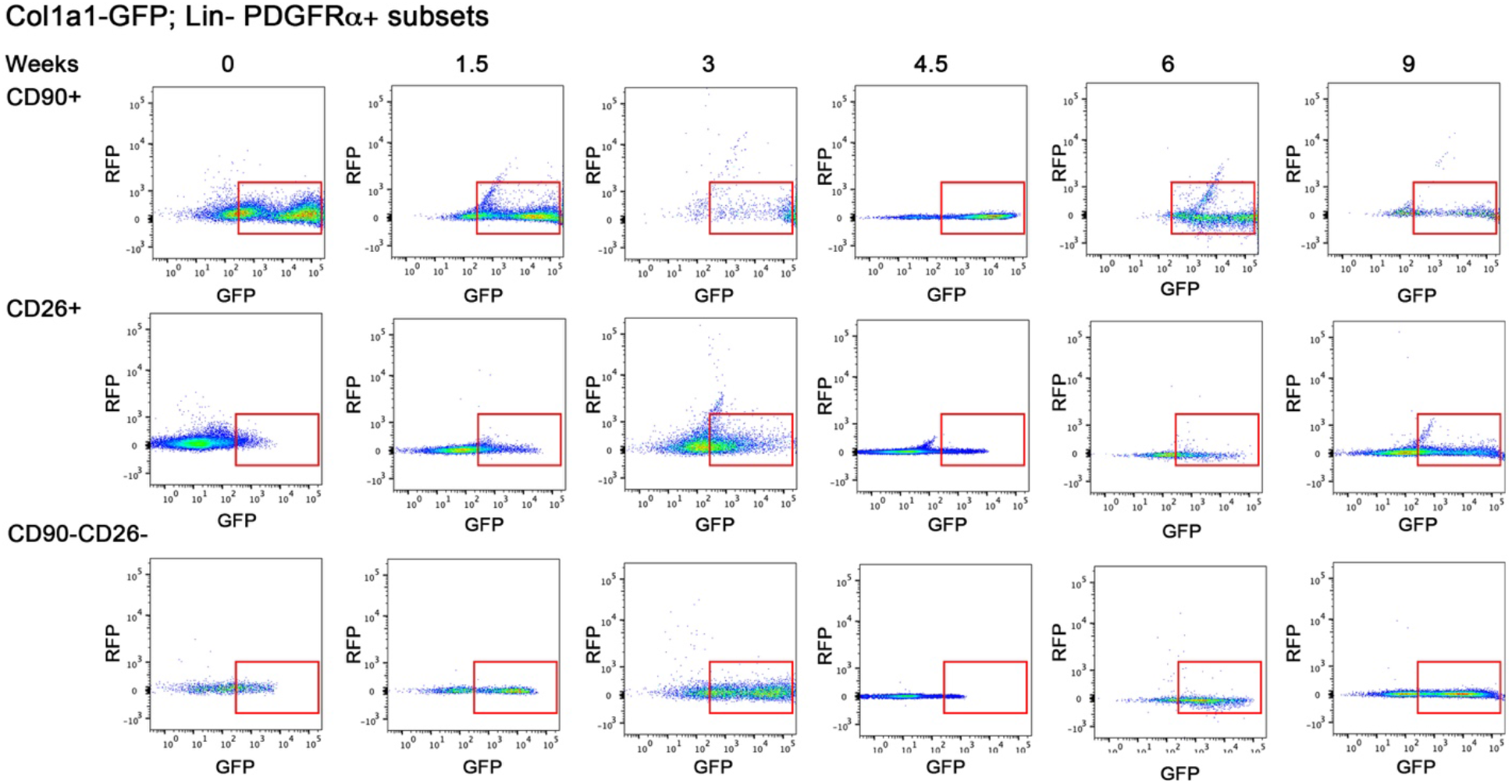
Col1a1 lineage expression of TdTomato to identify collagen expressing cells. Flow cytometry analysis of Col1-CreERT2;TdTomato mice treated with tamoxifen (Figure 2A) to identify Col1 expressing cells (TdTm+) over time after bleomycin. Representative flow plots of Lin(-) PDGFRα+ fibroblasts (upper panel) and Lin positive CD31, CD45 and EpCAM cells (lower panels).

**Supplementary Figure 6.**
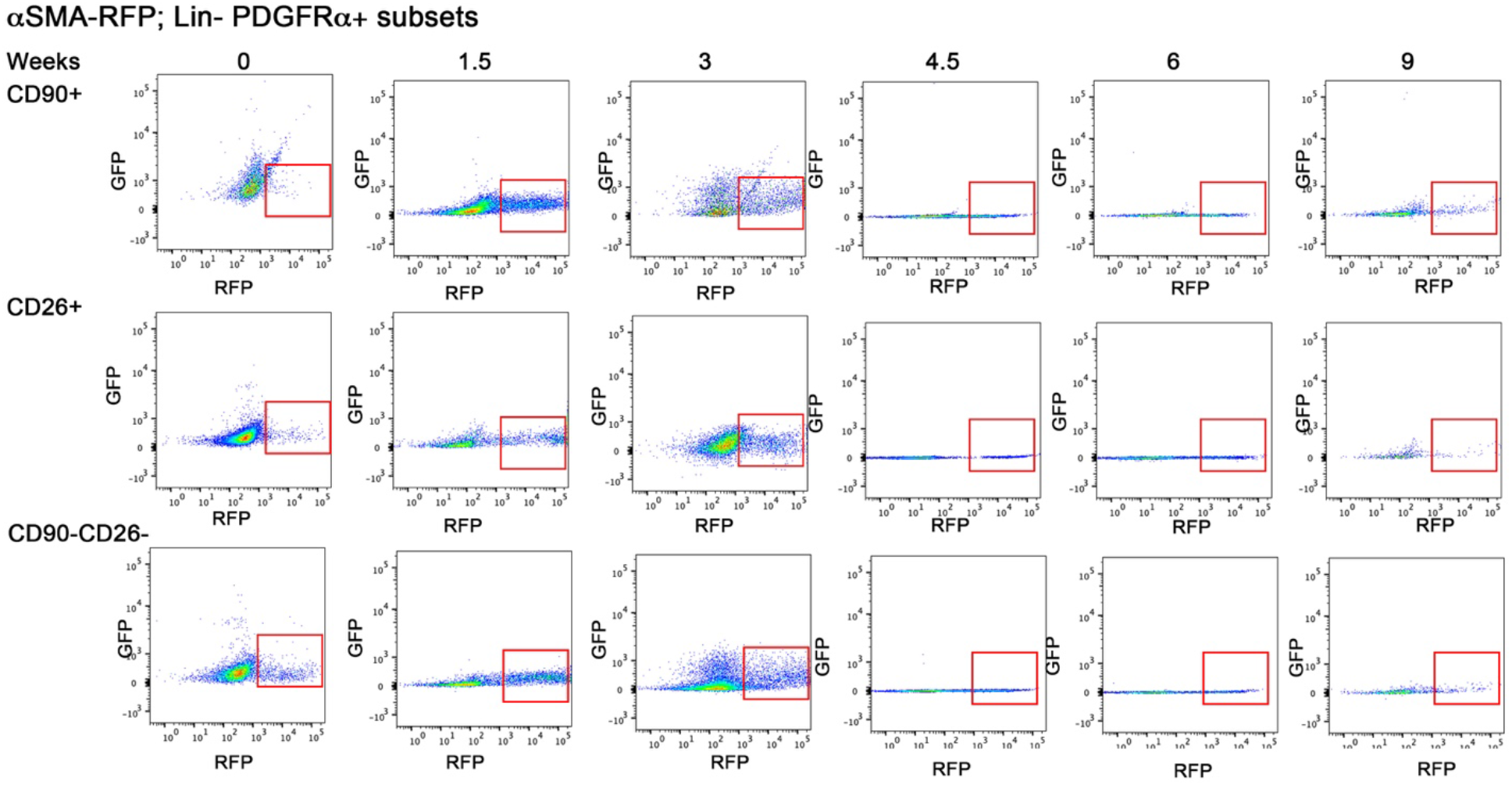
Col1-GFP expression of fibroblast subsets over time after bleomycin. Representative flow cytometry plots of GFP expression in CD90+CD26-, CD26+CD90- and CD90-CD26-fibroblasts subsets in Col-GFP mice during fibrosis development and resolution.

**Supplementary Figure 7.**
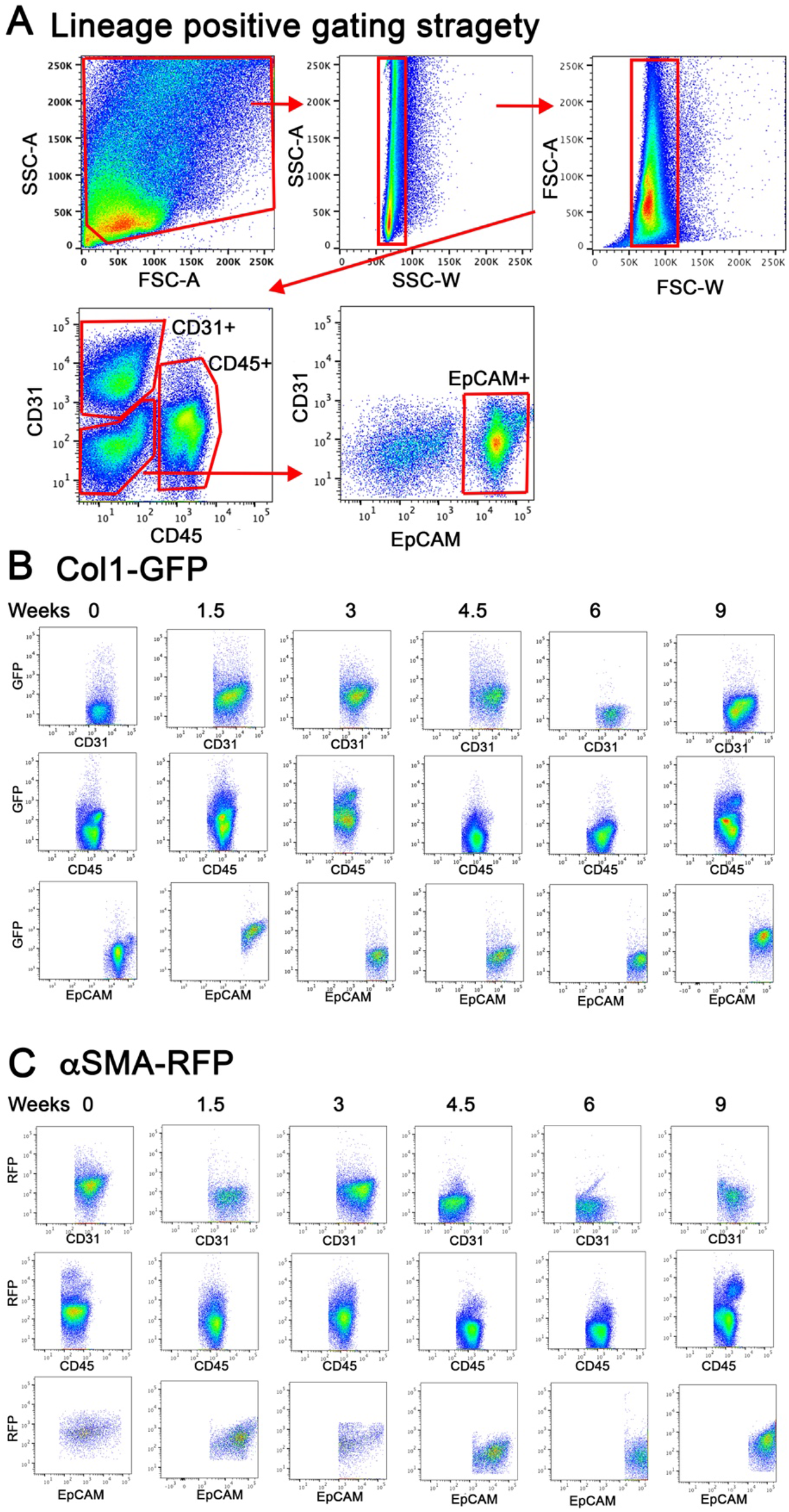
αSMA-RFP expression of fibroblast subsets over time after bleomycin. Representative flow cytometry plots of RFP expression in CD90+CD26-, CD26+CD90- and CD90-CD26-fibroblasts subsets in αSMA-RFP mice during fibrosis development and resolution.

**Supplementary Figure 8.**
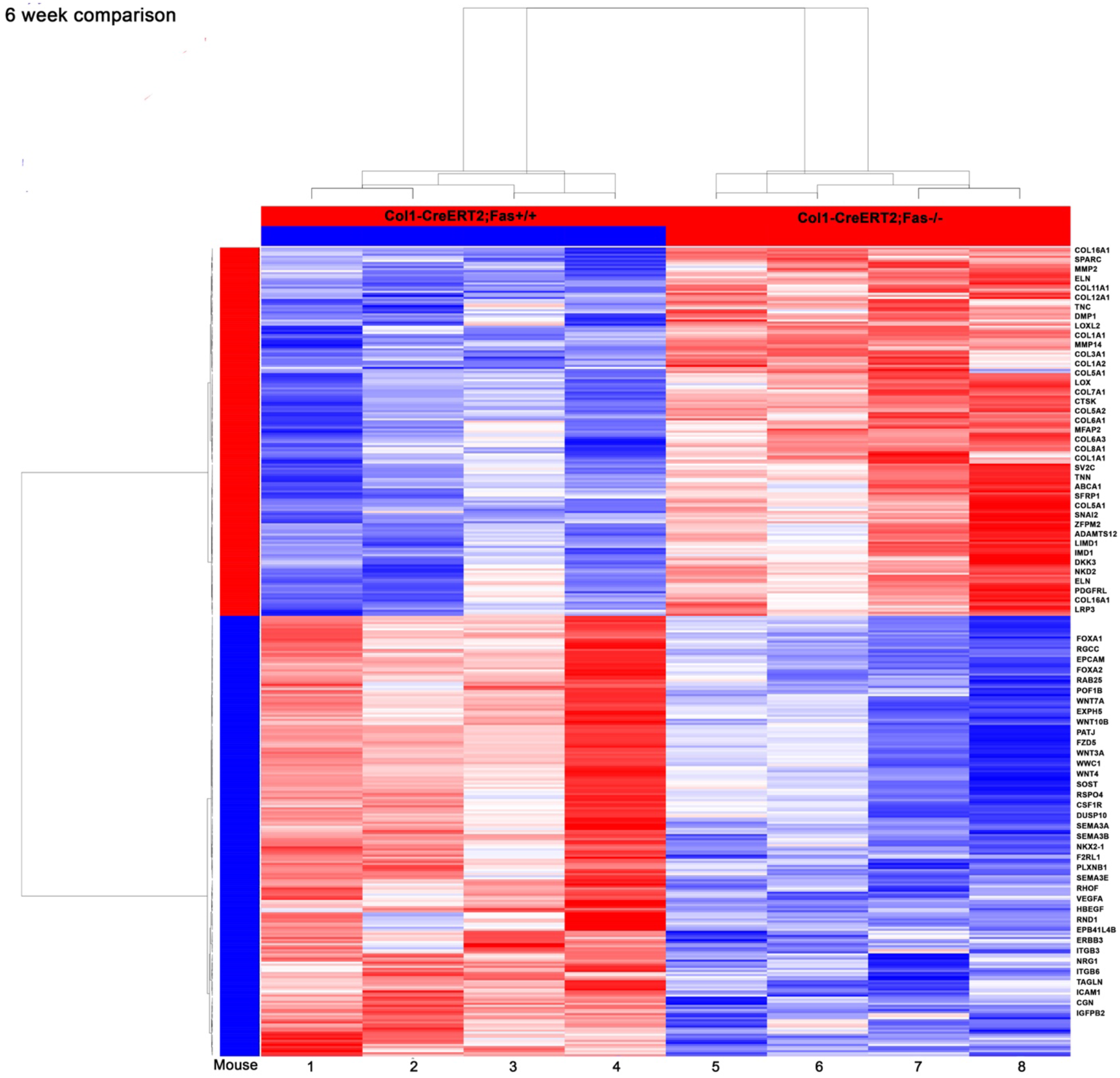
Transcriptome expression in bulk sequencing of Lin (-) fibroblasts. (A) Heat maps of normalized signal show transcriptional differences between Lin(-) fibroblasts from Col1-CreERT2;Fas^+/+^ and Col1-CreERT2;Fas^−/−^ mice at six weeks after bleomycin. (n=4/group)

**Supplementary Figure 9.**
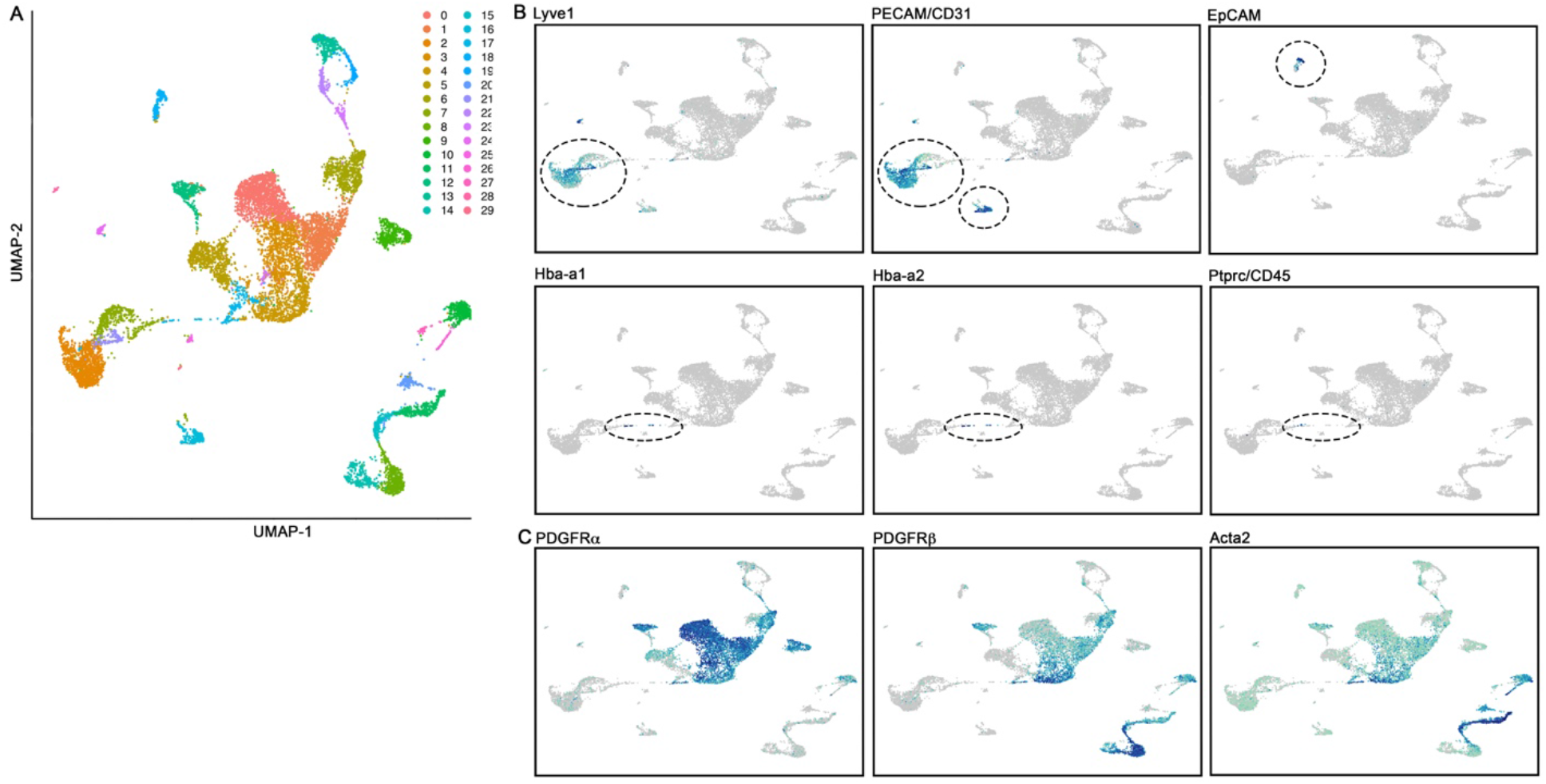
Cell clusters identified from Lin(-) sorting by sc-RNA sequencing. (A) UMAP of all samples pre-filtering. (B) UMAP overlays of gene expression identifying cell clusters as non-fibroblasts (C) UMAP overlays of gene expression identifying cell clusters as mesenchymal cells and fibroblasts.

**Supplementary Figure 10.**
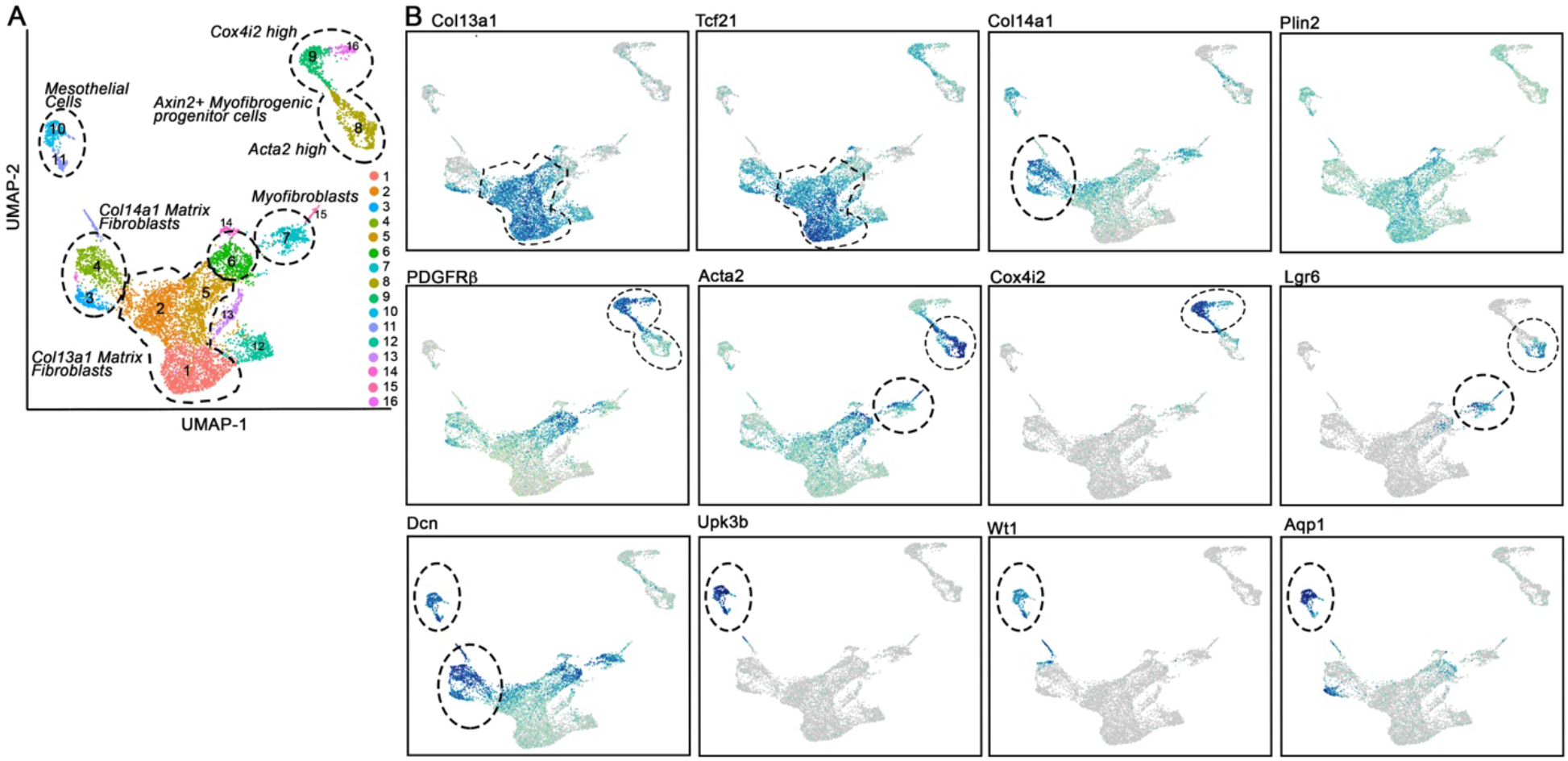
Sc-RNA cluster identification based from currently recognized populations. (A) UMAP of all samples with clusters identified/named from the literature (28,29). (B) UMAP overlays of gene expression identifying cell clusters.

**Supplementary Figure 11.**
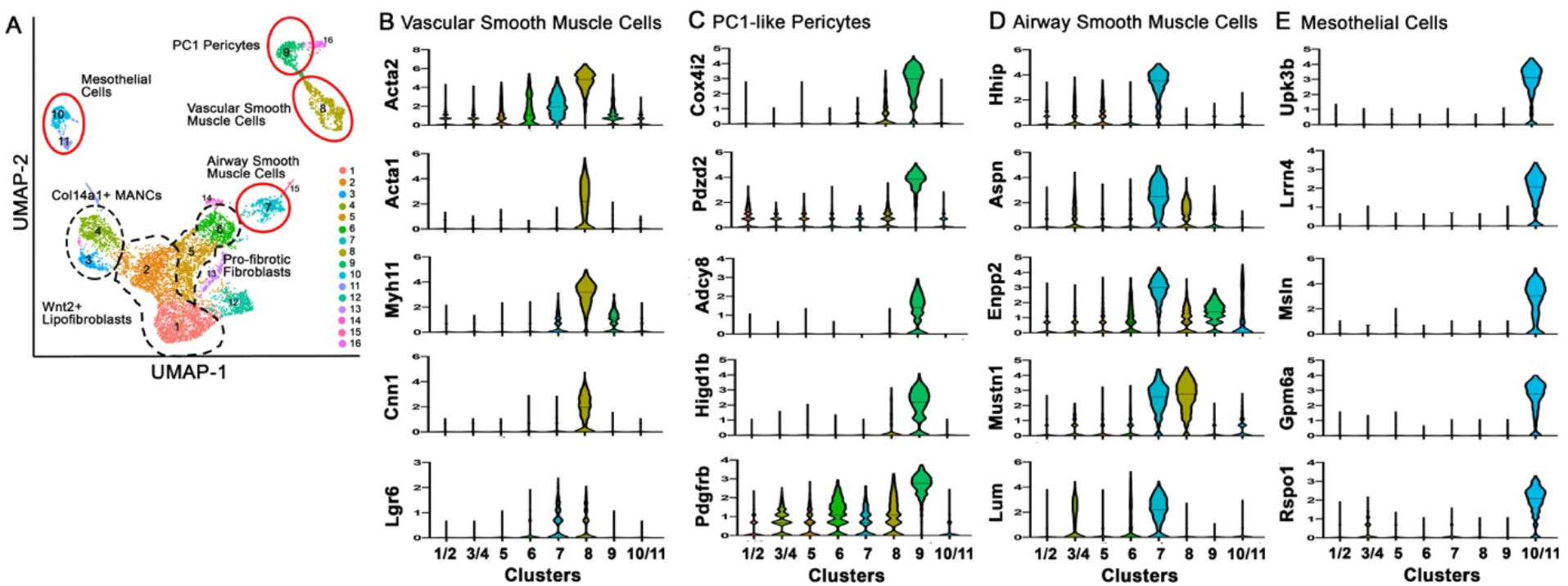
Sc-RNA sequencing cluster quality analysis. (A) RNA counts for identified clusters. (B) Feature RNA counts for identified clusters. (C) Percent ribosomal RNA counts for identified clusters. (D) Percent mitochondrial RNA counts for identified clusters. Cluster 12 was only distinguishable from cluster 1 due to low expression of ribosomal RNAs and high expression of mitochondrial RNAs. Cluster 13 was only differentiated due to its high levels of ribosomal RNAs. Cluster 15 was only distinguishable from cluster 7 due to high expression of mitochondrial RNAs. Cluster 16 was only distinguishable from cluster 9 due to1low expression of ribosomal RNAs and high expression of mitochondrial RNAs.

**Supplementary Figure 12.**
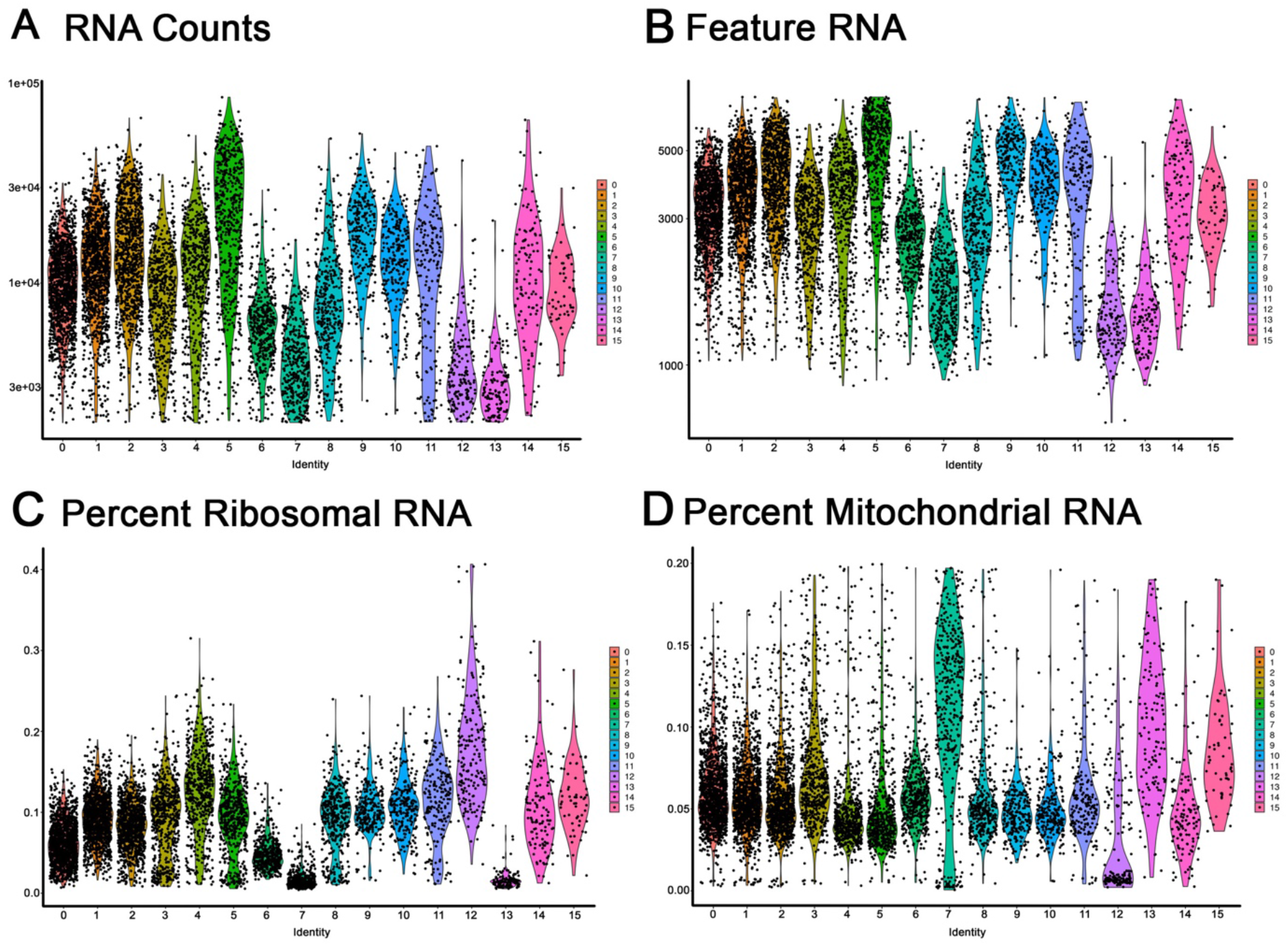
Gene expression profile of mesenchymal clusters. (A) UMAP of sc-RNA clusters. (B) Violin plots showing genes highly associated with vascular smooth muscle cells (cluster 8). (C) Violin plots showing genes highly associated with PC1-like pericytes (cluster 9). (D) Violin plots showing genes highly associated with airway smooth muscle cells (cluster7). (E) Violin plots showing genes highly associated with mesothelial cells (clusters 10 and 11).

**Supplementary Figure 13.**
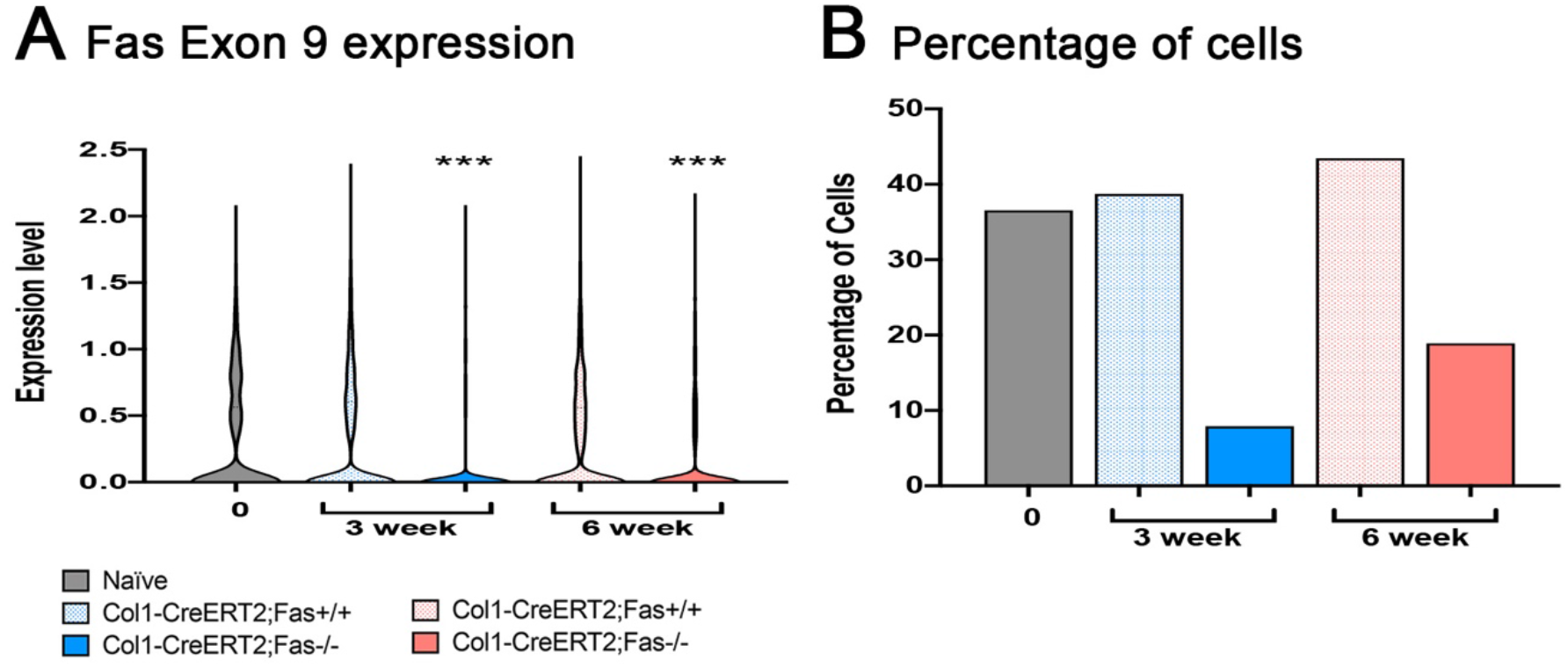
Expression of Fas exon 9. (A) Expression of Fas exon 9 in naïve, Fas sufficient and Fas-deficient cells. (B) Percentage of cells expression Fas exon 9 in naïve, Fas sufficient and Fas-deficient cells.

**Supplementary Table 1:** Enrichment pathways from bulk sequencing of Lin-fibroblasts at 6 weeks.

| **Genotype** | **Enrichment Pathways** | **Adjusted p value** |
| --- | --- | --- |
| Col1-CreERT2;Fas<sup>-/-</sup> | ECM organization | 6.32E-13 |
|  | Collagen fibril organization | 6.59E-09 |
|  | Endodermal cell differentiation | 1.45E-05 |
|  | Platelet-derived growth factor binding | 2.76E-05 |
|  | ECM receptor interactions | 0.00136 |
|  | Negative regulation of cell differentiation | 0.00618 |
|  | Negative regulation of canonical Wnt Signaling | 0.1489 |
| Col1-CreERT2;Fas<sup>+/+</sup> | Regulation of cell-cell adhesion mediated by cadherin | 0.00786 |
|  | Regulation of cell migration | 0.0079 |
|  | Epithelial cell development | 0.0357 |
|  | Hippo signaling | 0.0584 |
|  | Wnt signaling | 0.0622 |
|  | Wound healing | 0.0699 |

**Supplementary Table 2:** Enrichment pathways from single cell sequencing at 6 weeks between Col1-CreERT2;Fas^−/−^ and Col1Cre-ERT2;Fas^+/+^ mice.

| Cluster | Enrichment Pathways | Adjusted p value |
| --- | --- | --- |
| Col13a1 matrix fibroblasts (cluster 1) | Sulfur compound biosynthetic process | 0.000952 |
|  | Mesonephric tubule development | 0.000422 |
|  | Actin cytoskeleton reorganization | 0.00128 |
|  | Positive regulation of vasculature development | 0.00153 |
|  | Regulation of endothelial cell migration | 0.00208 |
| Col13a1 matrix fibroblasts (cluster 2) | Extracellular matrix organization | 0.00189 |
|  | Extracellular matrix assembly | 0.0175 |
|  | Endoplasmic reticulum lumen | 0.0057 |
|  | Elastic fibril formation | 0.0072 |
|  | Molecules associated with elastic fibers | 0.024 |
|  | Collagen biosynthesis and modifying enzymes | 0.0335 |
| Pro-fibrotic fibroblasts (cluster 5) | Extracellular matrix organization | 1.234E-15 |
|  | Regulation of cell migration | 3.662E-06 |
|  | Extracellular matrix disassembly | 1.963E-05 |
|  | Regulation of smooth muscle cell migration | 0.000851 |
|  | Cytokine-mediated signaling pathway | 0.000901 |
| Pro-fibrotic fibroblast (cluster 6) | Extracellular matrix organization | 1.916E-29 |
|  | Collagen fibril organization | 5.207E-10 |
|  | Skeletal system development | 7.199E-09 |
|  | Endoderm formation | 1.496E-07 |
|  | Regulation of cell migration | 5.711E-07 |
|  | Regulation of cell proliferation | 5.519E-07 |
|  | Negative regulation of apoptotic process | 3.599E-06 |
| Col14a1 matrix fibroblasts (cluster 3) | Co-translational protein targeting to membrane | 9.16E-20 |
|  | Cytosolic ribosome | 1.421E-18 |
|  | Protein targeting to ER | 1.999E-18 |
|  | Extracellular matrix organization | 9.297E-11 |
| Col14a1 matrix fibroblasts (cluster 4) | SRP-dependent co-translational protein targeting to membrane | 6.138E-88 |
|  | Co-translational protein targeting to membrane | 3.289E-86 |
|  | Protein targeting to ER | 4.807E-86 |
|  | Peptide biosynthetic process | 6.899E-63 |

**Supplementary Table 3:**
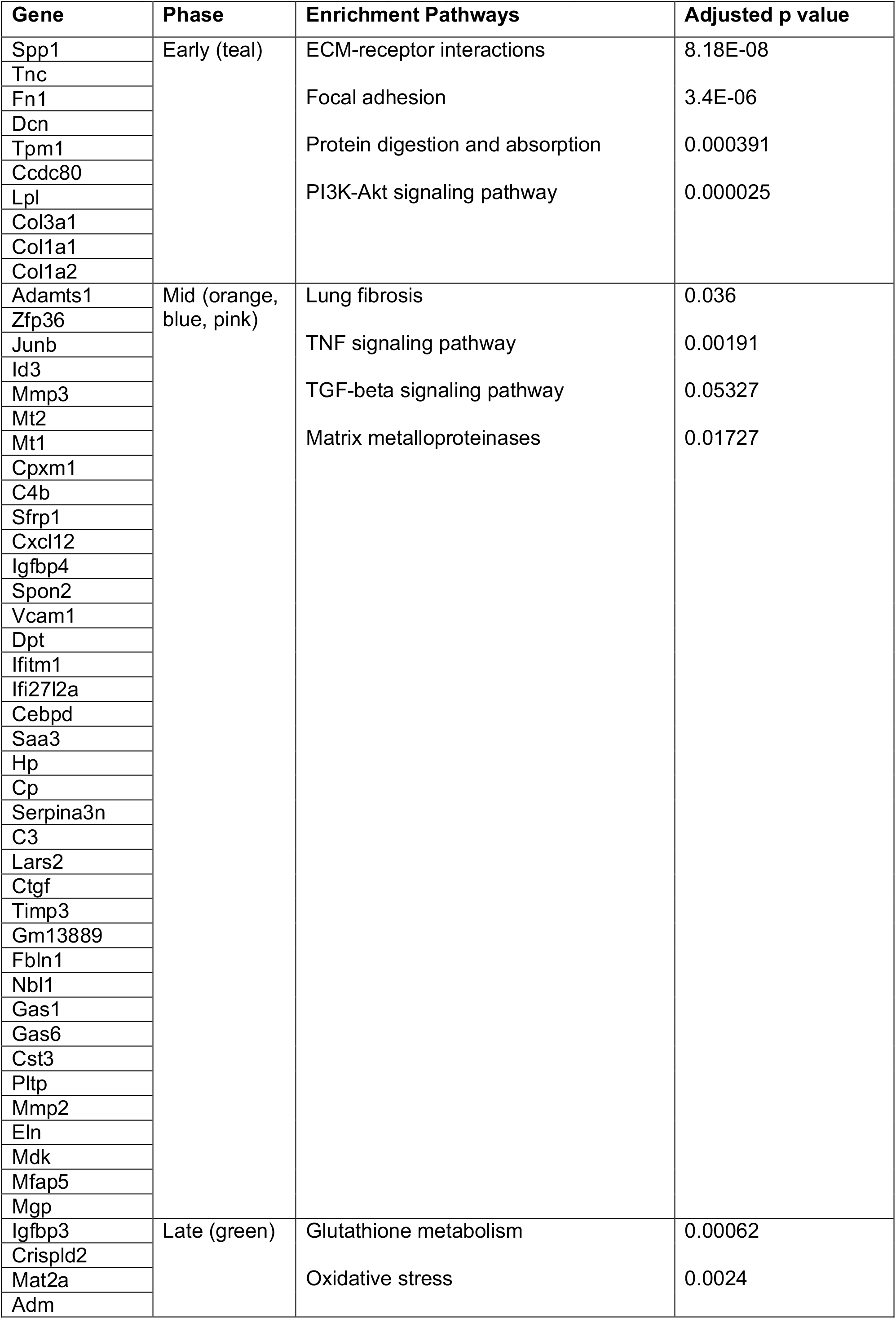
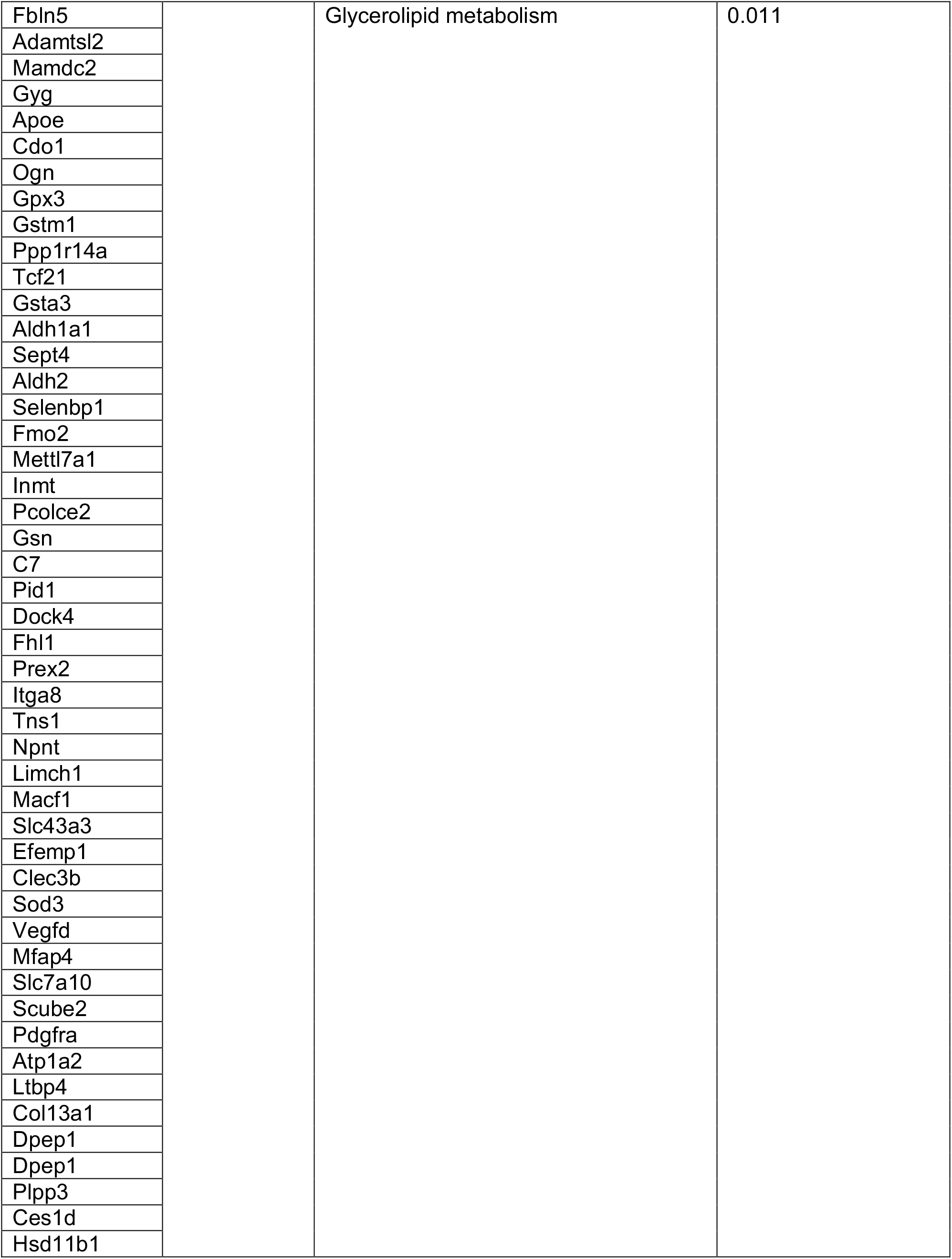
Pseudotime trajectory scenario 1 (clusters 1-5-6)

**Supplementary Table 4:** Psueotime trajectory scenario 2 (clusters 2-5-6)

| Gene | Phase | Enrichment Pathways | Adjusted p value |
| --- | --- | --- | --- |
| Thbs1 | Early<br>(orange) | ECM-receptor interactions | 2.03E-05 |
| Mt2 |  |  |  |
| Mt1 |  |  |  |
| Spp1 |  |  |  |
| Timp1 |  | Focal adhesion formation | 2.66E-05 |
| Tnc |  |  |  |
| Fn1 |  | PI3K-Akt signaling pathway | 0.000533 |
| Pltp |  |  |  |
| Mdk |  | Protein digestion and absorption | 0.000403 |
| Mfap5 |  |  |  |
| Pmepa1 |  |  |  |
| Eln |  |  |  |
| Lpl |  |  |  |
| Serpine2 |  |  |  |
| Mgp |  |  |  |
| Acta2 |  |  |  |
| Crip1 |  |  |  |
| Lgals1 |  |  |  |
| Dcn |  |  |  |
| Aebp1 |  |  |  |
| Igf1 |  |  |  |
| Fstl1 |  |  |  |
| Ccdc80 |  |  |  |
| Col3a1 |  |  |  |
| Col5a2 |  |  |  |
| Col1a1 |  |  |  |
| Sfrp1 | Mid (blue) | Cholesterol metabolism | 0.05256 |
| Igfbp4 |  | VEGF signaling pathway | 0.06193 |
| Saa3 |  |  |  |
| Hp |  | Glutathione metabolism | 0.06812 |
| Serpina3n |  |  |  |
| Cxcl12 |  |  |  |
| Gm13889 |  |  |  |
| Cxcl14 |  |  |  |
| Angptl4 |  |  |  |
| Ifi27l2a |  |  |  |
| Cebpd |  |  |  |
| Rgs2 |  |  |  |
| Hspb1 |  |  |  |
| Zfp36 |  |  |  |
| Id3 |  |  |  |
| Txnip |  |  |  |
| Adm |  |  |  |
| Wisp2 |  |  |  |
| Trf |  |  |  |
| Gyg |  |  |  |
| Cdo1 |  |  |  |
| Klf2 | Late (pink) | Matrix metalloproteinases | 0.08718 |
| Tsc22d3 |  | Pathways affecting insulin-like growth factor (IGF1)-Akt signaling | 0.08294 |
| Igfbp3 |  |  |  |
| Timp3 |  |  |  |
| Mmp3 |  |  |  |
| Spon2 |  |  |  |
| Igfbp6 |  |  |  |
| Nbl1 |  |  |  |
| Cpxm1 |  |  |  |
| C4b |  |  |  |
| C3 |  |  |  |
| Gas6 |  |  |  |
| Ogn |  |  |  |
| Apoe |  |  |  |
| Cst3 |  |  |  |
| Dpt |  |  |  |
| Efemp1 |  |  |  |
| Clec3b |  |  |  |
| Mamdc2 |  |  |  |
| Gpc3 |  |  |  |
| Vegfd |  |  |  |
| Sod3 |  |  |  |
| Mfap4 |  |  |  |
| Slc7a10 |  |  |  |
| Scube2 |  |  |  |
| Pdgfra |  |  |  |
| Ces1d |  |  |  |
| Aldh1a1 |  |  |  |
| Col13a1 |  |  |  |
| Hsd11b1 |  |  |  |
| Npnt |  |  |  |
| Atp1a2 |  |  |  |
| Prex2 |  |  |  |
| Itga8 |  |  |  |
| Tns1 |  |  |  |
| Limch1 |  |  |  |
| Macf1 |  |  |  |
| C7 |  |  |  |
| Gsta3 |  |  |  |
| Dpep1 |  |  |  |
| Plpp3 |  |  |  |
| Ppp1r14a |  |  |  |
| Tcf21 |  |  |  |
| Sept4 |  |  |  |
| G0s2 |  |  |  |
| Aldh2 |  |  |  |
| Mettl7a1 |  |  |  |
| Selenbp1 |  |  |  |
| Fmo2 |  |  |  |
| Pcolce2 |  |  |  |
| Inmt |  |  |  |
| Gsn |  |  |  |

